# A Hybrid Modeling Framework for Predictive Digital Twins of CHO Cell Culture

**DOI:** 10.1101/2025.11.24.690194

**Authors:** Anne Richelle, David Andersson, Athanasios Antonakoudis, Jesper Jakobsson, Shanti Pijeaud, Anton Vernersson, Johan Trygg

## Abstract

Digital twins of mammalian cell cultures hold great potential for predictive bioprocess modeling, yet their development is challenged by the nonlinear dynamics and metabolic complexity of these systems. We present a hybrid computational framework that integrates mechanistic and data-driven modeling to construct predictive digital twins for Chinese hamster ovary (CHO) cell cultures producing monoclonal antibodies. The framework couples ordinary differential equation (ODE) models with constraint-based metabolic modeling and machine learning components trained on Bayesian-estimated metabolic rates. Applied to 23 CHO fed-batch cultures, viable cell density, product titer, and key metabolite concentrations are accurately predicted under varying feeding and media conditions within a unified simulation engine, where empirical variability is incorporated through multivariate statistical constraints derived from experimental data. Cross-validation analyses demonstrated strong generalization across process variations, highlighting the framework’s capacity to capture both biochemical constraints and adaptive cellular behavior. This hybrid modeling approach provides a mechanistically interpretable yet data-adaptive foundation for constructing bioprocess digital twins. By bridging statistical, mechanistic, and machine learning methodologies, it advances the computational representation of CHO cell culture systems and offers a generalizable strategy for predictive modeling in complex biological production processes.

## 1. Introduction

Biopharmaceutical manufacturing increasingly relies on Chinese hamster ovary (CHO) cells as the primary host system for monoclonal antibody (mAb) production (Walsh and Walsh, 2022). While decades of process development have yielded robust and scalable fed-batch processes, cell culture performance remains highly variable due to the complex interplay of genetic, metabolic, and environmental factors. Achieving consistent outcomes requires a deep understanding of cellular metabolism and its response to dynamic culture environments, yet predictive capabilities remain limited. Traditional empirical strategies for process optimization are resource-intensive, while purely mechanistic modeling approaches often fail to capture the inherent complexity and variability of mammalian cell cultures. These challenges underscore the need for more integrative and predictive frameworks that can guide process development, optimization, and control in real time (Babi et al., 2022; Luo et al., 2021; Narayanan et al., 2022; Richelle et al., 2020).

Digital twins (i.e., virtual replicas of physical systems) have emerged as a transformative concept in bioprocessing (Portela et al., 2021; Smiatek et al., 2020). By integrating process knowledge with data-driven models, digital twins can continuously simulate and predict process behavior under varying conditions. This enables forecasting of critical quality attributes, proactive identification of suboptimal conditions, and rational design of feeding strategies and medium compositions. Despite their potential, digital twins for mammalian cell culture remain underexplored, largely due to the difficulty of integrating heterogeneous data sources with mechanistic knowledge in a scalable and predictive framework (Park et al., 2021; Park et al., 2024).

Several recent efforts highlight both the promise and limitations of current modeling approaches. Mechanistic and constraint-based frameworks have long been valued for their ability to provide insight into intracellular metabolism. For example, Yasemi and Jolicoeur (2023) introduced a genome-scale dynamic constraint-based modeling (gDCBM) framework for CHO cells, where uptake rates derived from exometabolite data were used to constrain fluxes in a dynamic flux balance analysis (dFBA) scheme. While this method enabled sequential predictions of intracellular fluxes and comparisons of metabolic states across CHO clones, it is inherently retrospective, requiring dense experimental data and lacking the ability to forecast or support process optimization.

Purely data-driven forecasting approaches are increasingly explored as a pathway toward digital twins. Park et al. (2023) developed multistep-ahead prediction models of mammalian cell culture performance, systematically evaluating combinations of forecasting strategies, AI architectures, and input features. Their models successfully predicted key performance indicators (including viable cell density (VCD), mAb titer, and concentrations of glucose, lactate, and ammonia) in near real-time. While powerful for monitoring and early-warning applications, these models offer limited mechanistic interpretability and cannot explicitly simulate the effects of novel interventions such as medium formulation or feed strategy changes.

Hybrid methods have sought to bridge these paradigms. Schinn et al. (2021) combined a genome-scale metabolic network with statistical learning to predict amino acid consumption in fed-batch CHO cultures. By coupling flux sampling with sigmoid function fitting, they were able to forecast amino acid depletion profiles, although the framework remained restricted to batch processes and could not accommodate altered feeding strategies. Ramos et al. (2022) proposed a hybrid semi-parametric flux balance model in which statistical regression parameterized selected fluxes within the network. While this improved flexibility relative to conventional FBA, the absence of dynamic integration across modeling layers limited its predictive generalizability.

Taken together, these studies underscore a common limitation: current methods are either retrospective, reliant on static fits, or unable to generalize predictions beyond narrowly defined scenarios. Reviews of the field (Park et al., 2021) highlight the transformative potential of digital twins to accelerate process development and enable adaptive control, but emphasize that most frameworks remain fragmented, focusing on retrospective analysis, narrow predictive tasks, or incomplete integration of mechanistic and data-driven models.

In summary, mechanistic models provide biological interpretability but struggle to forecast or adapt to new conditions; data-driven methods enable prediction but lack mechanistic grounding; and existing hybrid approaches remain static and loosely coupled. This landscape highlights the need for a truly integrative digital twin framework: one capable of combining mechanistic insight with data-driven adaptability, leveraging prior experimental knowledge, and dynamically simulating process trajectories.

In this work, we introduce such a hybrid digital twin framework for mammalian cell culture, validated using monoclonal antibody production in CHO cells. Our approach integrates multiple modeling paradigms (i.e., machine learning, mechanistic ordinary differential equation (ODE) models, constraint-based metabolic models, and statistical modeling) into a unified, automated simulation engine. Specifically, viable cell density (VCD) dynamics are predicted using machine learning models trained on metabolic rates estimated through a Bayesian approach (Andersson et al. 2025; Richelle et al., 2025), while dynamics of some predefined core metabolites (i.e., glucose, lactate, ammonium, glutamine, and glutamate) are captured by ODE-based kinetic rate models. These predictions are coupled to a reduced metabolic network model, dynamically constrained by prior experimental data through multivariate statistical analysis. To enable this integration, we build upon and extend the hybrid FBA model of Ramos et al. (2022) by removing static time dependencies, allowing dynamic forecasting of metabolite evolution from a reduced set of input variables.

The resulting digital twin framework enables predictive simulation of cell culture behavior under different feeding and media conditions, offering a versatile tool for process design, optimization, and control. Unlike existing retrospective or static approaches, our method supports forward-looking simulations that can guide decision-making during process development, reduce experimental burden, and enable adaptive process control. By bridging mechanistic insight with data-driven adaptability, this work provides a step toward operationalizing digital twins in the biopharmaceutical industry.

## 2. Materials and Methods

### 2.1. Experimental Dataset

The dataset consisted of 23 fed-batch Ambr® 15 mL bioreactor runs using CHO-S cells producing the therapeutic protein Xolair (Omalizumab). A daily sampling strategy was applied to monitor 25 metabolites, including viable cell density (VCD), product titer, glucose, lactate, ammonium, and 20 amino acids.

All batches were operated under the same process conditions, including inoculation and process control parameters, while variations were introduced at the level of feed composition and quantity. Three feed streams were employed: FMA, FMB, and FMG. FMA and FMB contained the full complement of carbohydrates, amino acids, and vitamins required for growth, but were separated into two feeds to avoid solubility issues at high concentrations. FMG consisted of a glucose stock solution that was manually added to maintain glucose levels above 5 g/L following bolus additions of FMA and FMB. The same FMG stock was used in all batches. Throughout the study, eight distinct FMA+FMB formulations were tested. For experimental sets with the same formulation, the volumes of FMA+FMB added varied, but the timing of additions was consistent across all experiments. The experimental data are detailed in Supplementary Figure 1.

### 2.2. Rate Calculation

Metabolic rates were inferred from concentration measurements using a Bayesian framework combined with nested sampling (MetRaC), as described previously in Andersson et al. (2025) and Richelle et al. (2025). The procedure begins by correcting raw metabolite profiles for reactor volume changes caused by feeding and sampling events. The resulting “pseudo-concentration” data are then fitted with a linear combination of basis logistic functions to capture consumption and production trends. Nested sampling–based Bayesian parameter estimation provides posterior distributions for the model parameters, thereby quantifying uncertainty and enabling systematic model comparison. This results in an ensemble of rate trajectories representing the full posterior distribution. Rates can subsequently be extracted at arbitrary time points, increasing the effective temporal resolution of the experimental data while retaining uncertainty information. In the present study, 100 trajectories were computed per metabolite and per culture, each evaluated at evenly spaced time points (apart from the training of the neural network for prediction of VCD dynamics, for which 200 trajectories were sampled per metabolite and per culture). For each time point, the mean as well as the 5th and 95th percentiles of the posterior rate distribution were calculated. The framework can also be used to back-propagate the pseudo-concentration posterior to reconstruct the concentrations at any sampling time points and capture the variability “perceived” by the method.

### 2.3. Neural Network for VCD prediction

The viable cell density (VCD) predictor was implemented as a dense neural network with a recurrent structure to capture temporal dependencies in the data (Mienye, et al. 2024). Input features include specific consumption rates and extracellular metabolite concentrations derived from the MetRaC framework (Section 2.2). The model is designed to predict the time-resolved specific growth rate (*μ* (*t*)) by combining a baseline estimation with a deviation correction mechanism:

- Baseline Growth Rate Estimation: A sigmoid-based regression model interpolates the mean specific growth rate across all training batches (i.e., 21 batches across all feeding conditions randomly selected out of the 23 available batches). This baseline represents the average growth behavior of the dataset, independent of specific input features.
- Deviation Prediction: A neural network predicts deviations from this mean growth profile based on input data. This component enables the model to tailor predictions to each culture condition, capturing dynamic responses to changes in metabolite availability.

The final predicted growth rate is obtained by adding the predicted deviation to the baseline growth rate.

The predictor can be expressed as :

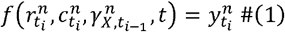

and optimized by minimizing the loss:

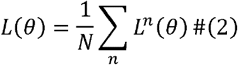

where

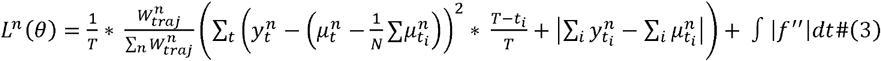

and

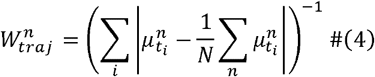

Here *μ* represent ground truth specific growth rates, and y represent the corresponding predictions of the model. *r* and *c* represent the specific consumption rates and the metabolite concentrations, respectively. The index *n* indexes trajectories, *i* indexes time points within a trajectory, *T* is the total number of trajectories, *T* is the batch end time (i.e., the maximum time), each batch contains 200 trajectories.

The first loss term penalizes deviations from the mean growth rate 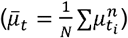. It is implemented as a mean squared error (MSE) between the predicted and ground-truth mean and is evaluated across all trajectories of all batches. Two weighting schemes shape this term: trajectory weights and time weights. Trajectory weights 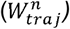, applied in both the first and second loss terms, upweight trajectories whose mean specific growth rate is near the batch mean and downweight outliers (e.g., atypical MetRaC decreasing profile such as 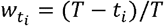, which increases the influence of early timepoints. trajectories). Time weights, applied only to the first term, emphasize early dynamics using a Highlighting early behavior improves convergence and predictive performance because early growth strongly determines later productivity. Downweighting outlier trajectories prevents the model from overfitting noise and keeps training focused on representative batch dynamics. The second term constrains the average predicted trajectory to match the ground truth mean, improving end-point VCD predictions. The final smoothness term penalizes sharp fluctuations by integrating the absolute second derivative of the predictions made (computed numerically via *torch*.*diff(n=2)* along the time dimension), encouraging biologically realistic dynamics.

Recurrence is introduced through the computation of the “biomaterial” variable, defined as:

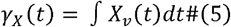

where *X*_*V*_ (*t*) is the viable cell density. Using the growth rate relation 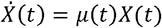, the discrete updates become:

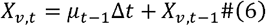

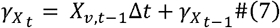

The model predicts deviations in growth rate, and consequently, deviations in biomaterial accumulation. The biomaterial term captures latent biological effects not directly observed in the data, such as accumulated inhibitory by-products (Richelle et al., 2022).

At each time step *t*_*i*_, the network receives the specific consumption rates 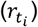, metabolite concentrations 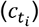, and the biomaterial value 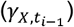 associated with the previously predicted growth rate 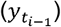. These inputs are concatenated and passed through a series of linear layers that expand and contract the feature space (Supplementary Figure 2).

With 21 metabolites as inputs (i.e., all metabolites except glucose, lactate, and ammonium), the initial feature dimension is 45 (21 rates + 21 concentrations + previous growth rate + time + timestep). This is projected to 64 dimensions, processed through three hidden layers, and then reduced to a single output neuron, the number of neurons in each layer is 64, 64, 64, 16, 8, 1. Each hidden layer uses GeLU activation and layer normalization, with dropout applied between layers for regularization.

Finally, the mean rate interpolator, composed of four fitted sigmoid functions, provides the baseline prediction that is adjusted by the deviation network output to yield the final specific growth rate prediction.

### 2.4. ODE Kinetic Models for Biomass Population and FLEX Metabolites

We developed a kinetic model comprising two interconnected components: an ODE-based biomass population kinetic model and an ODE-based metabolite model. The latter describes five key metabolites (i.e., glucose, lactate, glutamine, ammonium, and glutamate) referred to as “FLEX” metabolites, as they are measured using the BioProfile® FLEX2 analyzer from Nova® Biomedical.

#### 2.4.1. ODE Biomass Populations Kinetic model

The ODE-based biomass population kinetic model was developed to estimate the effective death rate of the culture. This step is essential because the growth rate calculated directly from viable cell density (VCD) data, denoted as *μ*, represents only the net rate of change in viable biomass over time. However, to properly constrain the metabolic model that will be used further in this study, we require the effective growth rate (*μ*_*eff*_), which reflects the actual rate of biomass synthesis, independent of cell death and lysis.

In this framework, the CHO cell population is divided into three subgroups: viable cells (*X*_*v*_), dead cells (*X*_*d*_), and lysed cells (*X*_*l*_). Cell death and lysis are driven by the accumulation of metabolic byproducts, represented by a catch-all “biomaterial” variable, *γ*_*X*_.

Following the approach of Richelle et al. (2022), the dynamics of these cell populations and of *γ*_*X*_ are described by the following system of ordinary differential equations :

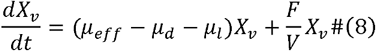

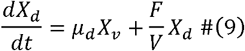

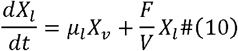

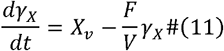

where *F*/*V* is the dilution rate, *μ*_*eff*_ is the effective growth rate, *μ*_*d*_ the death rate, and *μ*_*l*_ the lysis rate.

In this study, we assumed that dead (*X*_*d*_) and lysed (*X*_*l*_) cells are a direct degradation product of the viable cell population (*X*_*v*_) (Kroll et al., 2016). However, since the analytical assay used population (*X*_*l*_) was maintained as an internal model variable rather than an experimentally (trypan blue exclusion) does not distinguish between dead and lysed cells, the lysed cell constrained quantity.

The effective death and lysis rates were modeled as base rates (*k*_*d*_ and *k*_*l*_) modulated by toxicity factors (*k*_*Td*_ and *k*_*Tl*_) linked to the accumulation of biomaterial or lysed cells, respectively:

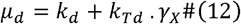

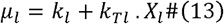

Finally, the effective growth rate (*μ*_*eff*_) can be derived from the net VCD growth rate (*μ*) calculated via MetRaC (Section 2.2) as follows:

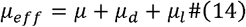

This model, therefore, provides an estimate of *μ*_*eff*_, necessary for accurate metabolic model constraints, by accounting for the contributions of both cell death and lysis that are not captured in *μ* alone.

#### 2.4.2. ODE FLEX Metabolites Model

An ordinary differential equation (ODE) model was developed to describe the dynamics of five key extracellular metabolites, referred to as “FLEX” metabolites: glucose (*Glc*), lactate (*Lac*), glutamine (*Gln*), glutamate (*Glu*), ammonia (*NH*4). The system consists of six state variables (including the viable cell concentration, *X*_*v*_) and captures both cellular metabolism and reactor-level dynamics (e.g., feeding and dilution). The governing equations are given in Eqs. (15–19):

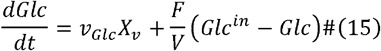

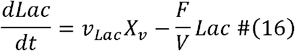

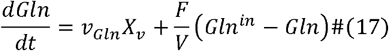

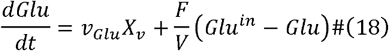

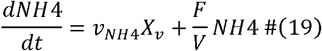

where *v*_*Glc*_, *v*_*Lac*_, *v*_*Gln*_, *v*_*Glu*_ and *v*_*NH*4_ are the specific consumptions and/or production rates. *Glc*^*in*^, *Gln*^*in*^ and *Glu*^*in*^ are metabolite concentrations in the feed. *F* is the feeding rate and *V* is the bioreactor volume.

The glucose consumption rate (*v*_*Glc*_) accounts for substrate saturation, inhibition by lactate, and maintenance energy requirements. It is proportional to the effective growth rate (*μ*_*eff*_), as shown in Eq. (20):

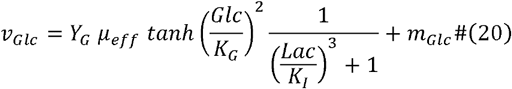

where *Y*_*G*_ is a pseudo-stoichiometric yield coefficient (glucose consumed per unit of effective growth rate), *K*_*G*_ is the half-saturation constant for glucose, *K*_*I*_ the inhibition constant for lactate, and *m*_*Glc*_ the maintenance coefficient for glucose.

Lactate metabolism was modeled in the context of the overflow metabolism (Lao and Toth, 1997), a phenomenon where cells excrete partially oxidized byproducts despite aerobic conditions. This occurs when the glucose uptake rate exceeds the oxidative capacity, leading to lactate accumulation. The lactate production (*v*_*Lac,Prod*_) and consumption (*v*_*Lac,Cons*_) rates are defined as:

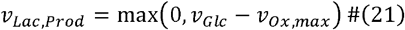

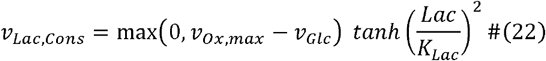

where *v*_*Ox,max*_ is the maximum oxidative capacity and *K*_*Lac*_ the half-saturation constant for lactate reuptake. The net lactate flux is then computed as:

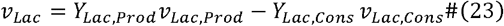

where *Y*_*Lac,Prod*_ and *Y*_*Lac,Cons*_ are yield coefficients representing the balance between production and consumption phases.

The glutamate consumption rate (*v*_*Glu*_) follows a similar formulation to glucose, scaled by the effective growth rate and accounting for maintenance energy:

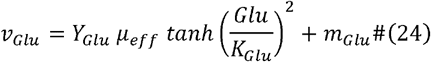

where *Y*_*Glu*_ is the yield coefficient, *K*_*Glu*_ the half-saturation constant for glutamate, and *m*_*Glu*_ the maintenance coefficient for glutamate.

The consumption of glutamine (*v*_*Gln*_) is modelled as :

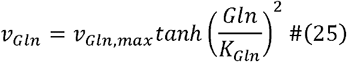

where *v*_*Gln,max*_ represents the maximum uptake rate, and *K*_*Gln*_ the half-saturation constant.

Ammonium production (*v*_*NH*4_) is modeled as a byproduct of glutamate and glutamine catabolism:

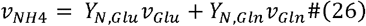

where *Y*_*N,Glu*_ and *Y*_*N,Gln*_ are the yield coefficients for ammonium formation from glutamate and glutamine, respectively.

### 2.5. Metabolic Model Construction

A stepwise model reduction pipeline was applied to derive a minimal, yet biologically and computationally consistent, metabolic model of CHO cells (Antonakoudis and Richelle, 2025). This approach ensures that the reduced model remains both feasible and predictive at each stage of the reduction process. Starting from the genome-scale reconstruction iCHO1766 (Hefzi et al., 2016; also referred to as iCHOv1), the reduction was performed iteratively by removing non-essential reactions while systematically verifying model feasibility.

Briefly, the reduction procedure involved (i) resolving infeasibilities, (ii) identifying essential exchanges required for model solvability, (iii) pruning redundant transport and inactive reactions based on parsimonious flux balance analysis (pFBA), and (iv) eliminating thermodynamically infeasible cycles. Dead-end metabolites and blocked reactions arising during each step were identified and removed to maintain network consistency. The pipeline was implemented using COBRApy (Ebrahim et al., 2013).

Experimental flux constraints were applied using the 95% confidence interval of metabolic rates estimated by MetRaC. Three demand reactions were added to ensure feasibility (step 1 of the reduction process): tyrosine demand (−0.00736 mM/gDCW.h ≤ v ≤ 0), lysine demand (−0.00750 mM/gDCW.h ≤ v ≤ 0), and threonine demand (−7×10^−^□ mM/gDCW.h ≤ v ≤ 0). Additionally, cystine exchange (EX_Lcystin_e_, present in the feed but not measured) was retained for feasibility (step 2 of the reduction process).

The final reduced model reproduces all expected auxotrophies except for asparagine. The asparagine synthase reaction (ASNS1) was initially removed during pFBA due to zero flux across all conditions, likely reflecting sufficient extracellular asparagine availability. To ensure biological completeness, ASNS1 was reintroduced and fixed in the final model. The evolution of model composition and the characteristics of the final reduced network are summarized in Supplementary Table 1.

### 2.6. PC-dFBA algorithm

To model dynamic metabolic flux distributions, we developed a hybrid semi-parametric modeling approach termed Principal Component–dynamic Flux Balance Analysis (PC-dFBA). PC-dFBA builds upon the HybridFBA formulation originally proposed by Ramos et al. (2022), which combines mechanistic (“parametric”) constraints from Flux Balance Analysis (FBA) with empirical (“non-parametric”) constraints derived from multivariate data analysis. The evolution of the algorithm, from HybridFBA through its dynamic and machine-learning–based refinements, is summarized in Supplementary Table 2 and detailed in the Supplementary Methods.

In its final form, PC-dFBA integrates four key components:

1. Mechanistic constraints: stoichiometric mass balance, reaction directionality, and measured exchange flux bounds, as in standard FBA.
2. Empirical constraints: flux-correlation relationships captured through Principal Component Analysis (PCA)–derived equations.
3. Dynamic simulation: sequential optimization across time, ensuring smooth temporal transitions via a MOMA (Minimization of Metabolic Adjustment; Segrè et al., 2002) continuity constraint.
4. Machine-learning–based parameterization: an artificial neural network (ANN) model predicts PCA loadings directly from exchange rates, replacing static PCA decompositions and removing the dependency on fixed time intervals.

The hybrid linear program simultaneously optimizes intracellular and exchange fluxes (*v*) and PCA scores (*Score*) :

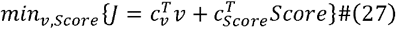

subject to:

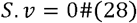

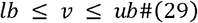

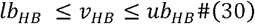

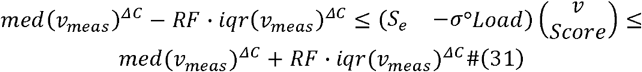

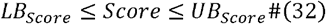

In this formulation, *S* denotes the stoichiometric matrix of the metabolic network, and *v* represents the vector of reaction fluxes, including both intracellular and exchange fluxes. The subset of exchange fluxes constrained by hard bounds (*v*_*HB*_) corresponds to the metabolites used as inputs in the simulation (i.e., defined by the 5 FLEX metabolites and the VCD for this study). Their lower and upper limits, *lb*_*HB*_ and *ub*_*HB*_, are fixed to the mean exchange rates for the simulated batch obtained using MetRaC. When imposing these bounds makes the linear program infeasible, they are relaxed to the 5^th^ and 95^th^ percentiles of the corresponding rate distributions. *S*_*e*_ is the stoichiometric matrix associated with the extracellular metabolites for which exchange rates are measured. The vector *c*_*v*_ specifies the FBA objective, typically the biomass growth reaction, while *c*_*Score*_ is a vector of ones that minimizes the empirical (PCA-related) contribution to the objective function.

The term *RF* represents a relaxation factor that allows limited deviation between predicted and measured fluxes, while *med* and *iqr* refer to the median and interquartile range (IQRs) of the measured exchange rates computed with MetRaC. The use of medians and IQRs, rather than means and standard deviations as in Ramos et al. (2022), provides greater robustness to experimental variability and outliers; the IQR, defined as the difference between the 75^th^ and 25^th^ percentiles, better captures the typical range of observed flux behavior. The lower and upper bounds of the PCA scores, *LB*_*Score*_ and *UB*_*Score*_, restrict the feasible range of each principal component.

To account for variation in data reliability, the culture duration was divided into two intervals *ΔC*: from day 0 to day 3, and from day 3 until the end of the culture. This separation was necessary because the metabolic rate estimates generated by MetRaC during the initial accuracy. Finally, the term *σ*°*Load* represents the normalized PCA loadings predicted directly phase (days 0–3) exhibited higher uncertainty and otherwise led to degraded prediction from exchange rate inputs by an artificial neural network (ANN), eliminating explicit dependence on time-windowed PCA of the original algorithm version and allowing continuous, data-driven estimation of empirical constraints.

To develop and train the ANN, time-series metabolite exchange rates from fed-batch CHO cultures were first discretized into 0.5-day windows. For each window, a PCA was performed using only batches grown under identical media and feeding conditions. This window-specific, condition-matched decomposition yielded biologically consistent “ideal” PCA loadings, resulting in one to five PCs capturing ≥90% of the variance. Both the 0.5-day window size and the 90% variance threshold were selected after testing multiple alternatives (results not shown), as these choices consistently produced the lowest overall SSE between model predictions and experimental data. These loadings served as target values for training the artificial neural network (ANN) models, each of which was designed to model a single principal component, thereby providing a continuous mapping between metabolite exchange fluxes and their corresponding PCA loadings. This strategy replaces static, time-dependent PCA decompositions with a flexible, data-driven representation capable of generalizing across batches and culture conditions.

To ensure consistency across datasets, exchange rates (features *X*) and corresponding scaled PCA loadings (*σ*°*Load*, targets *Y*) were concatenated across all time intervals and batches, grouped by PC (PC1, PC2, …). Missing PCs were replaced with zero-filled vectors matching the feature dimension to maintain a fixed output dimensionality (≤5 PCs explaining ≥85–90% of the variance). All inputs and targets were standardized (zero mean, unit variance) using *StandardScaler* from *Scikit-learn*, and the scalers fitted on the training data were reused to normalize new inputs and de-normalize model predictions.

The ANN training followed a two-step procedure. In the first step, hyperparameters were optimized using the *Optuna* framework by minimizing the root mean squared error 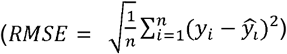 between observed and predicted PCA loadings, while monitoring mean absolute error (MAE) and R^2^ scores. The optimized parameters included the number of hidden layers, neurons per layer, batch size, learning rate, L2 regularization strength, and dropout rate. Hyperparameter optimization was conducted separately for two model configurations: *Model_All*, trained on all metabolites as features, and *Model_Flex*, trained only on the subset of FLEX metabolites together with viable cell density (VCD). During this optimization, the dataset was assumed to satisfy the IID (independent and identically distributed) condition, meaning that all samples were treated as independent and drawn from the same distribution. Accordingly, the data were randomly shuffled and split into training, validation, and test subsets in a 70/15/15 ratio.

In the second step, the best hyperparameters identified by *Optuna* were reused to retrain the network architectures under different validation strategies to evaluate generalization. Three strategies were considered: (i) direct validation (IID setting), in which the ANN was trained on all media groups and tested on randomly held-out samples not used during training; (ii) leave-one-media-group-out (LOMO), in which all batches from one media group were excluded from training and used solely for testing, allowing assessment of the model’s ability to generalize to unseen media formulations; and (iii) leave-one-batch-out (LOBO), where each batch was sequentially excluded from training and used for testing as an independent validation case. Preliminary analyses (data not shown) indicated that the optimal hyperparameters were largely consistent across media groups; therefore, the same architectures and training configurations obtained during the initial optimization were applied to all validation strategies to ensure a fair comparison of predictive performance.

At runtime, the PC-dFBA algorithm operates as a sequential dynamic simulation, iterating over discrete time intervals (0.1 days in this study). Each iteration involves four main computational steps:

1. **Input processing and normalization**. At each time point, measured or estimated extracellular exchange rates are provided as input to the model. These rates are first normalized using the same *StandardScaler* parameters (mean and standard deviation) that were applied during ANN training, ensuring consistency between simulation inputs and the data distribution used for model calibration.
2. **Prediction of principal component loadings**. The normalized exchange-rate vector one of the five principal components (PCs). Each ANN predicts a *σ* · *Load* vector is passed through the trained artificial neural networks (ANNs), each corresponding to representing the loadings associated with its respective PC. The predicted loadings are then de-normalized and assembled into a complete loading matrix, which captures the empirical relationships among fluxes at that time step.
3. **Flux optimization under hybrid constraints**. The hybrid linear program (LP) is then formulated and solved to determine intracellular and exchange fluxes. The predicted *σ*°*Load* matrix replaces static PCA terms in the empirical constraint set, while mechanistic constraints are imposed on mass balance, flux bounds, and the subset of hard bounds exchange rates referred to as hard bounds. The LP thus combines mechanistic realism with data-driven flexibility, producing flux distributions consistent with both metabolic stoichiometry and empirical flux correlations.
4. **Dynamic update and flux continuity**. Once the optimization is solved, extracellular metabolite concentrations are updated using the computed fluxes to advance the simulation to the next time interval. Exchange fluxes that are not constrained as hard bounds are dynamically predicted by the PC-dFBA algorithm at each iteration. To ensure smooth temporal transitions and prevent abrupt, non-physical changes in flux profiles, a MOMA-style penalty (Segrè et al., 2002) minimizes the deviation of flux values between successive time points:

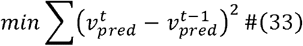 For the initial time step, where no prior flux distribution is available, standard FBA with PCA-derived constraints is used as the baseline solution.

In this study, the PC-dFBA framework was trained on metabolic rate data for 25 extracellular metabolites measured across 23 batches. During simulation, six metabolites, corresponding to the growth rate and exchanges associated with the FLEX metabolites, were specified as fixed hard bounds, while the remaining 19 exchange fluxes were dynamically predicted by PC-dFBA. This setup allowed the model to infer unmeasured fluxes, capture realistic metabolic correlations, and reproduce the variability observed across experimental conditions.

## 3. Results

### 3.1. General overview of the modeling framework for cell culture simulation

We developed a digital twin framework that integrates viable cell density (VCD) modeling using a recurrent neural network, kinetic modeling of extracellular metabolites, and constraint-based metabolic modeling into a unified platform for bioprocess simulation. The framework predicts the evolution of key culture variables, such as VCD, product titer, and metabolite concentrations, under any candidate culture medium and feeding strategy, based on prior experimental data and cell line–specific characteristics. By coupling mechanistic and data-driven components, the model captures both the biochemical constraints of the system and the empirical variability observed across bioprocesses.

The framework combines three core modeling components: a cell-specific metabolic model constrained using data-driven constraints, an ODE-based kinetic model describing the dynamics of key extracellular metabolites, and a neural network model predicting viable cell density. These components are integrated into a bioreactor simulator that dynamically computes fluxes and mass balances, thus enabling realistic simulation of process evolution over time. Two main data and information sources serve as inputs to the framework: (i) a genome-scale metabolic network describing all known metabolic reactions for the host cell, and (ii) a database of experimental data comprising process metadata, measured metabolite concentrations, and feeding profiles from previous bioprocesses using the same cell line.

The digital twin construction follows an automated workflow (Figure 1). First, experimental concentration data are converted into metabolic rates using the MetRaC (Metabolic Rate Calculation) approach, which applies a Bayesian estimation and nested sampling method to obtain robust rate estimates. These rates serve as the foundation for both kinetic model calibration and data-driven constraint identification. Simultaneously, the genome-scale metabolic network is refined to build a cell-specific model representing the actual metabolism of the host cell line. This reduction incorporates biological knowledge and observed exchange fluxes to tailor the network to the production context.

**Figure 1.**
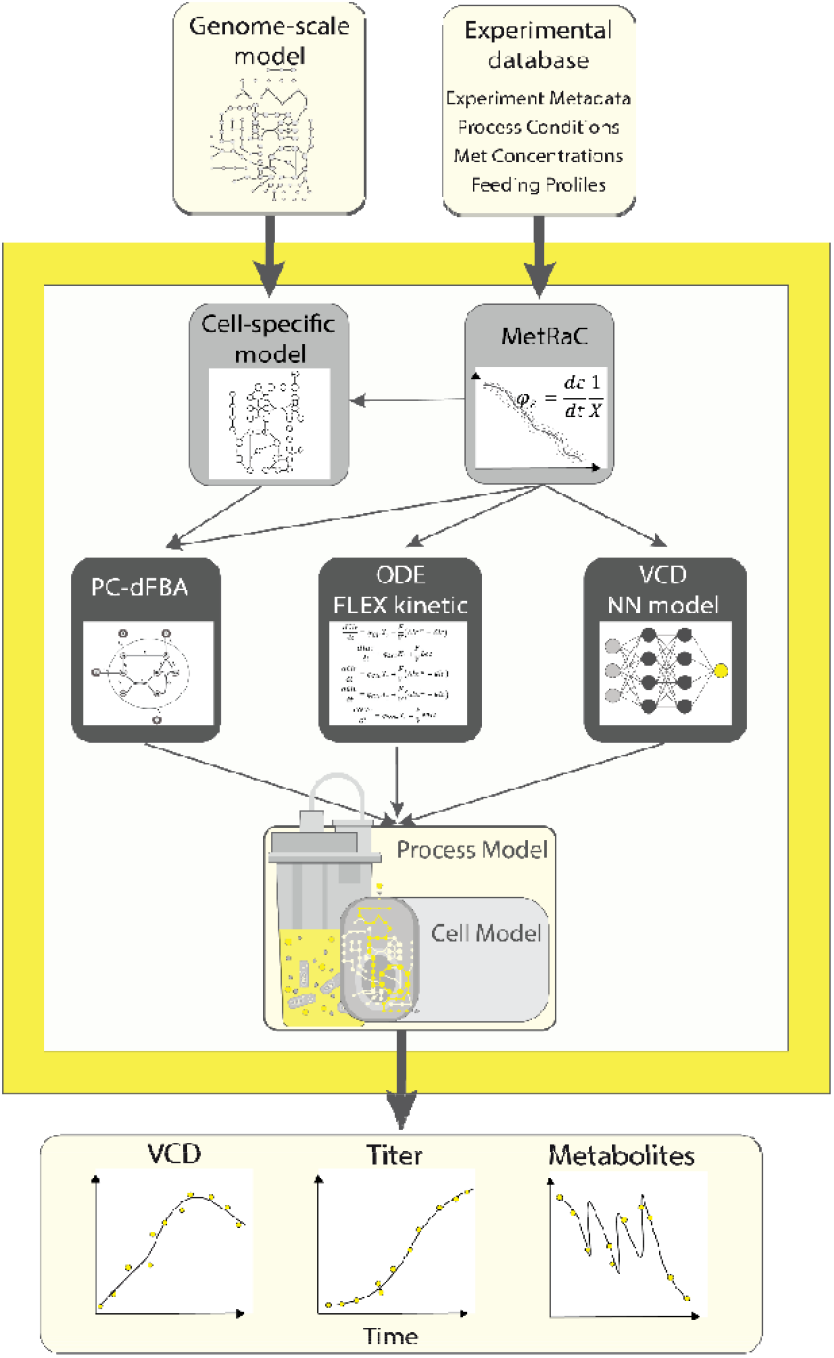
Schematic overview of the cell culture digital twin framework. The workflow integrates two main data and information inputs (i.e., a genome-scale metabolic model and an experimental database of process data) to construct a cell-specific model and calculate metabolic rates using the MetRaC approach. These data feed three modeling components: (i) the PC-dFBA metabolic model, (ii) the ODE-based FLEX kinetic model, and (iii) the VCD neural network model. The components are combined within a hybrid bioreactor simulator that dynamically predicts viable cell density, product titer, and metabolite concentrations over time.

Three modeling layers are then trained or parameterized based on the available processed data. The VCD neural network model learns to predict viable cell density trajectories from process data, capturing nonlinear growth behavior. The FLEX kinetic model, a reduced system of ODEs, describes the dynamics of key extracellular metabolites such as glucose, lactate, ammonia, glutamine, and glutamate. In parallel, multivariate data analysis is used to identify empirical relationships among metabolic fluxes and metabolite evolution, which are encoded as additional constraints in the Principal Component–dynamic Flux Balance Analysis (PC-dFBA) formulation. This hybrid FBA model integrates the mechanistic mass-balance constraints of metabolism with the statistical dependencies derived from experimental data, improving predictive power across culture conditions.

Finally, the three modeling components are integrated within a bioreactor simulator that performs dynamic mass balance calculations according to the general formulation:

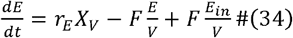

where *E* represents the extracellular metabolite concentration, *r*_*E*_ the rate predicted by either the VCD, kinetic, or metabolic model, *X*_*V*_ the viable cell density, *F* the feeding rate, *V* the bioreactor volume, and *E*_*in*_ the concentration of *E* in the feed. The resulting simulation environment enables prediction of the complete culture trajectory over time, including viable cell growth, product accumulation, and metabolite consumption or secretion.This digital twin framework provides a flexible and extensible foundation for in-silico process simulation. It can replicate experimental processes using measured feeding strategies or explore new hypothetical conditions by modifying input profiles.

### 3.2. Neural network-based prediction of VCD dynamics

To enable predictive modeling of cell growth within the digital twin framework, a neural network (NN) was trained to estimate the growth rate (, net rate of change in viable biomass over time) based on extracellular metabolite concentrations and specific consumption rates derived from the MetRaC pipeline (see Section 2.3 for more details). The dataset used for model development consisted of 200 trajectories sampled from the MetRaC model fitted for each of the variables, each corresponding to time-resolved rate and concentration profiles representative of the experimental variability observed across bioreactor runs. Out of the available datasets, 91% (21 batches) were used for training and 9% (2 batches) were withheld for validation.

The model was trained for 250 epochs using a batch-wise sequential strategy: during each epoch, trajectories from individual bioreactor batches were presented in random order. For each batch, the NN predicted the specific growth rate trajectory *μ* (*t*), computed the loss, and performed backpropagation before proceeding to the next batch. The training loss combined three complementary terms that jointly enforce accuracy, stability, and biological realism. Specifically, the loss penalized (i) the mean squared error between predicted and experimental growth rate trajectories, weighted by both trajectory representativeness and time point importance; (ii) deviations between the average predicted and target growth rates across trajectories, ensuring consistency in overall growth behavior; and (iii) excessive curvature of the predicted trajectory through a smoothness regularization term proportional to the integrated absolute second derivative.

To avoid overfitting and improve temporal generalization, the number of time points per trajectory was limited to 32, and 200 trajectories were used. This constraint reduced the effective recurrence depth and helped the model focus on meaningful time-dependent trends rather than stochastic fluctuations. Importantly, because the model learns from the cumulative biomaterial function rather than absolute timestamps, it generalizes across different sampling frequencies and process time scales, as long as the underlying biological trajectory remains comparable to those seen during training.

The trained NN accurately reproduced the specific growth rate profiles across all process conditions (Figure 2), capturing both the characteristic exponential growth and subsequent decline phases. When coupled with the ODE-based biomass balance equation, the predicted growth rates were translated into viable cell density (VCD) trajectories that closely matched experimental data (Supp Figure 3). The integration of the NN growth predictor into the ODE framework thus enables dynamic simulation of biomass accumulation and decline, forming a key component of the cell-process digital twin.

**Figure 2.**
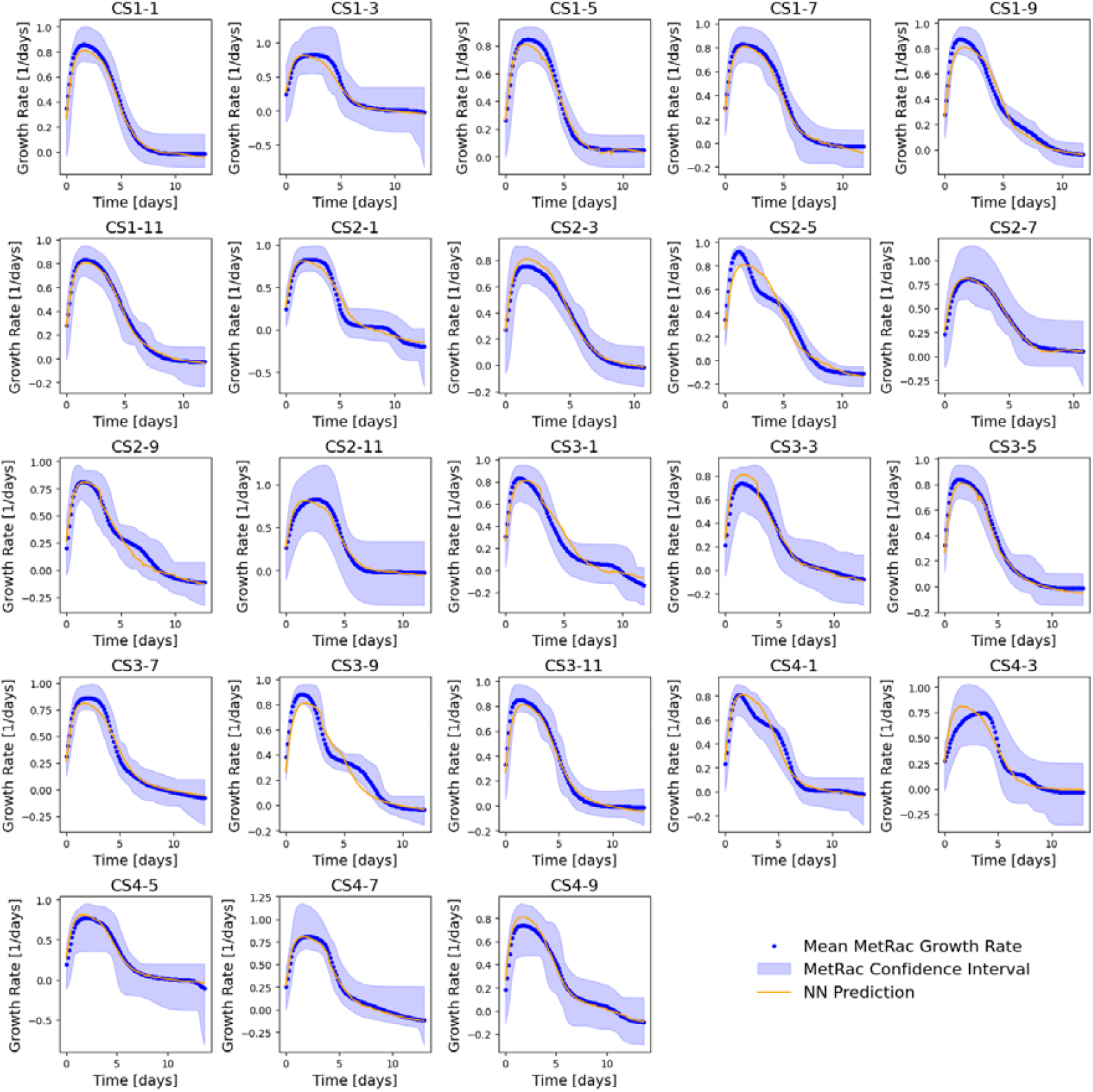
Predicted and experimental specific growth rate trajectories across batches. The blue shaded areas represent the MetRac confidence intervals, blue dots correspond to mean MetRaC growth rates, and the orange lines show model predictions.

### 3.3. ODEs model identification

The ODE-based kinetic modeling framework consisted of two interconnected modules: a biomass population model and a FLEX metabolite model (Section 2.4). These components were successively identified and validated using experimental data from 23 fed-batch cultures.

The first step involved estimating the effective growth rate (*μ*_*eff*_) using the ODE-based biomass population kinetic model. Model parameters were identified using the Nelder–Mead simplex algorithm to minimize the least-squares criterion between simulated and experimental viable and dead biomass profiles. To ensure biological realism, a constraint was introduced to maintain comparable magnitudes between dead and lysed cell pools, preventing divergence during parameter optimization while retaining model flexibility.

The resulting *μ*_*eff*_ trajectories reproduced the expected biological dynamics of CHO cell cultures, showing an initial exponential growth phase followed by a gradual decline during the stationary and death phases. Small negative *μ*_*eff*_ values occasionally appeared under low cell activity; these were truncated to 1×10^−^□ day^−1^ to maintain numerical and biological consistency.

A comparison between model-estimated *μ*_*eff*_ and experimentally derived *μ* values (computed from viable cell density via MetRaC) for all batches is presented in Figure 3. The identified parameter values are summarized in Table 1, and a detailed view of the fitted model performance, including viable, dead, lysed, and biomaterial profiles for each batch, is provided in Supplementary Figure 4. The model successfully captured the main growth and death dynamics across all cultures, validating its suitability as the foundation for the subsequent metabolite model.

**Table 1.**
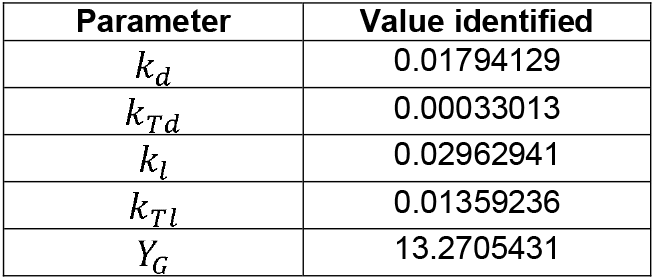
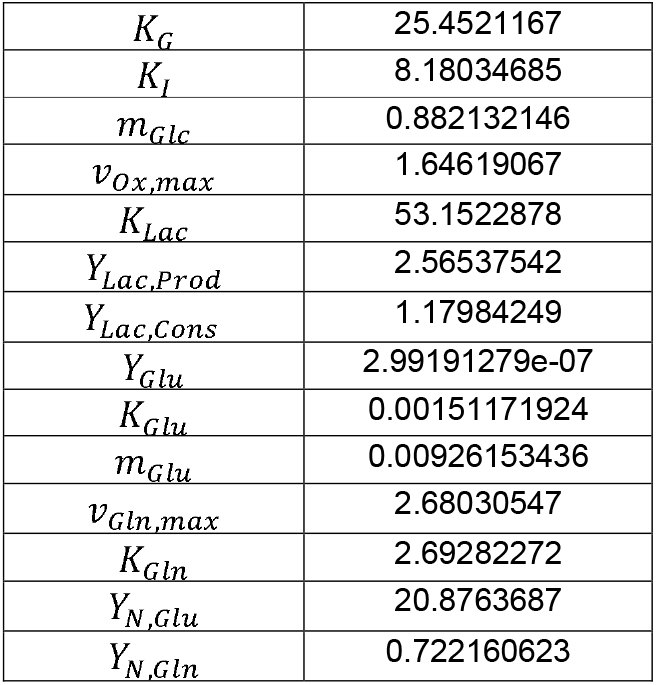
Parameter value identified for the ODEs biomass and FLEX metabolite models.

| Parameter | Value identified |
| --- | --- |
| $k_d$ | 0.01794129 |
| $k_{Td}$ | 0.00033013 |
| $k_l$ | 0.02962941 |
| $k_{Tl}$ | 0.01359236 |
| $Y_G$ | 13.2705431 |
| $K_G$ | 25.4521167 |
| $K_I$ | 8.18034685 |
| $m_{Glc}$ | 0.882132146 |
| $v_{Ox,max}$ | 1.64619067 |
| $K_{Lac}$ | 53.1522878 |
| $Y_{Lac,Prod}$ | 2.56537542 |
| $Y_{Lac,Cons}$ | 1.17984249 |
| $Y_{Glu}$ | 2.99191279e-07 |
| $K_{Glu}$ | 0.00151171924 |
| $m_{Glu}$ | 0.00926153436 |
| $v_{Gln,max}$ | 2.68030547 |
| $K_{Gln}$ | 2.69282272 |
| $Y_{N,Glu}$ | 20.8763687 |
| $Y_{N,Gln}$ | 0.722160623 |

**Figure 3.**
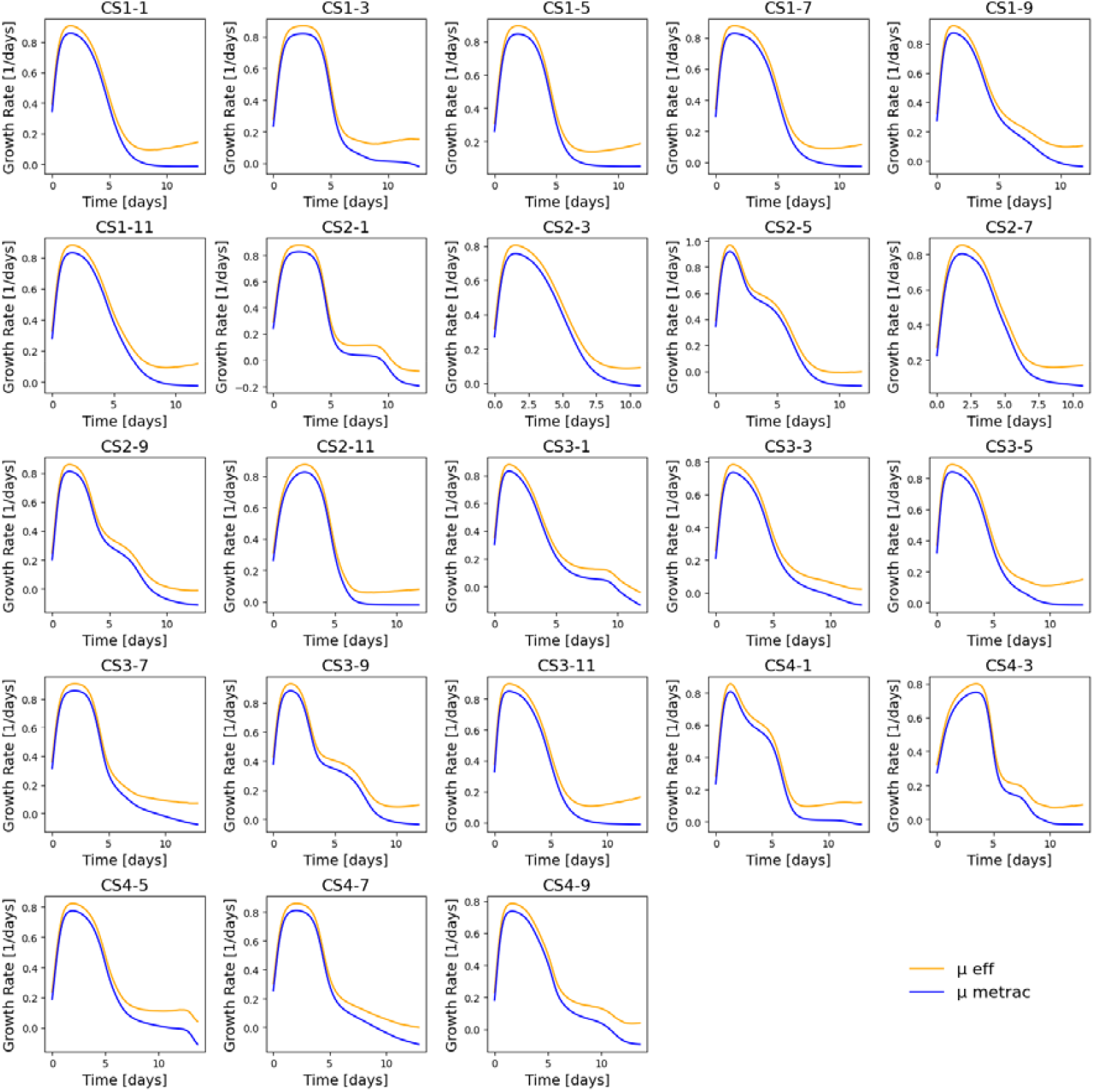
Comparison between the effective growth rate estimated by the ODE model (*μ*_*eff*_, orange) and the experimental growth rate derived from viable cell density data (*μ*, blue) for each of the 23 batches.

In the second step, the identified *μ*_*eff*_ trajectories were used to impose the viable cell dynamics (*X*_*v*_) in the ODE-based FLEX metabolite model. The model parameters were then optimized using the Nelder–Mead simplex algorithm to minimize the sum of squared differences between simulated and measured concentrations of five extracellular metabolites: glucose, lactate, glutamine, glutamate, and ammonia (Table 1).

The resulting model fits are summarized in Figure 4, which compares predicted versus measured metabolite concentrations across all batches, along with the corresponding coefficients of determination (R^2^). The model achieved good agreement for most metabolites, with R^2^ values of 0.91 for glucose, 0.98 for glutamine, 0.91 for glutamate, and 0.92 for ammonia, indicating accurate reproduction of the main consumption and production dynamics.

**Figure 4.**
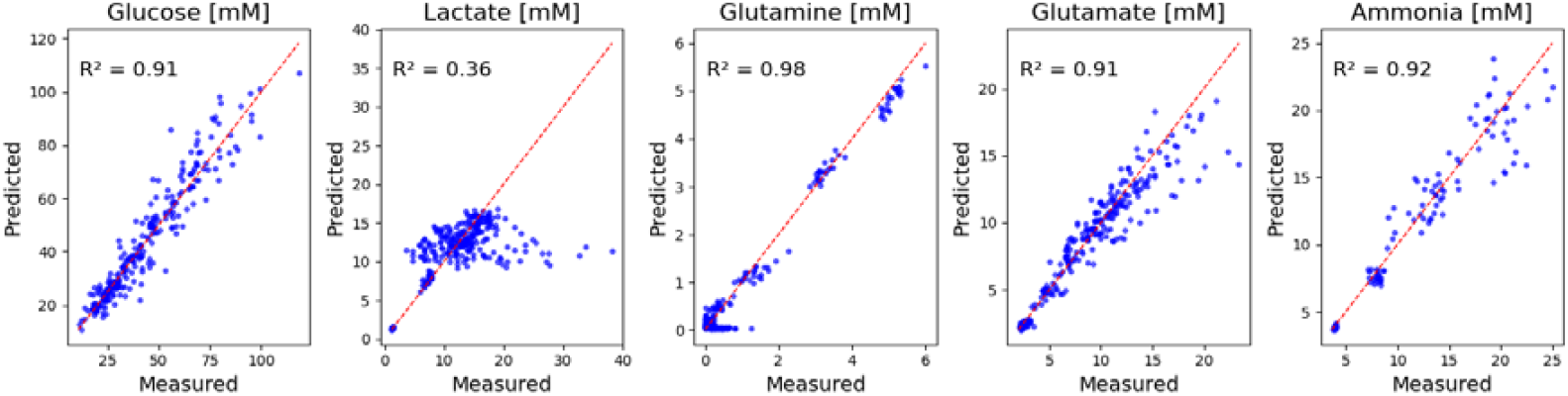
Direct validation of the FLEX ODE metabolite model. Scatter plots of predicted versus measured metabolite concentrations for glucose, lactate, glutamine, glutamate, and ammonia across all 23 experimental batches. Red dashed lines indicate the line of equality. The coefficient of determination (R^2^) for each metabolite quantifies the overall fit quality.

However, lactate exhibited a lower R^2^ of 0.36, reflecting the model’s limited ability to capture lactate re-accumulation observed during the later culture stages. This discrepancy is attributed to the simplified formulation of the current model, which includes only a small subset of extracellular metabolites. In particular, the lack of other metabolites involved in energy and redox balance (e.g., alanine, pyruvate, or TCA intermediates) restricts the model’s ability to reproduce lactate production shifts under metabolic stress. Extending the model to include these additional metabolic branches could improve its ability to describe late-phase lactate behavior, but is beyond the scope of the present study.

A detailed view of metabolite concentration profiles over time for all 23 experimental batches is provided in Supplementary Figure 5. Overall, the FLEX ODE model captures the main trends and magnitudes of metabolite dynamics, supporting its use as a mechanistic link to constrain subsequent metabolic modeling steps.

### 3.4. PC-dFBA training and predictions

The predictive performance of the PC-dFBA algorithm was evaluated using two ANN-based loading-prediction model configuration (i.e., *Model_Flex* and *Model_All*) combined with three validation strategies: direct validation, leave-one-media-group-out (LOMO), and leave-one-batch-out (LOBO). Direct validation corresponds to an IID setting in which all media compositions are represented during training, whereas LOMO and LOBO assess the model’s ability to generalize to unseen media compositions and individual batches, respectively (see Section 2.6. for details)

Model performance was quantified using the coefficient of determination (R^2^) and the summed squared error (SSE) for each metabolite concentration profile across 19 simulated extracellular metabolites, excluding the hard bounds metabolites with trivial identity mapping (glucose, lactate, glutamine, glutamate, ammonium, and VCD). Figure 5A summarizes the R^2^ scores across the 19 predicted metabolites and the six “*ANN x validation*” type combinations. Overall, PC-dFBA achieved strong predictive performance, with median R^2^ values ranging from 0.6 to 0.9 for 10 metabolites. The highest accuracies were obtained for branched-chain and aromatic amino acids (isoleucine, leucine, valine, phenylalanine, and threonine), which consistently reached R^2^ > 0.8 across all validation strategies. In contrast, alanine, asparagine, glycine, proline, and tyrosine displayed lower R^2^ values, reflecting either greater biological variability or a weaker mapping between their exchange fluxes and PCA-derived latent dynamics.

**Figure 5.**
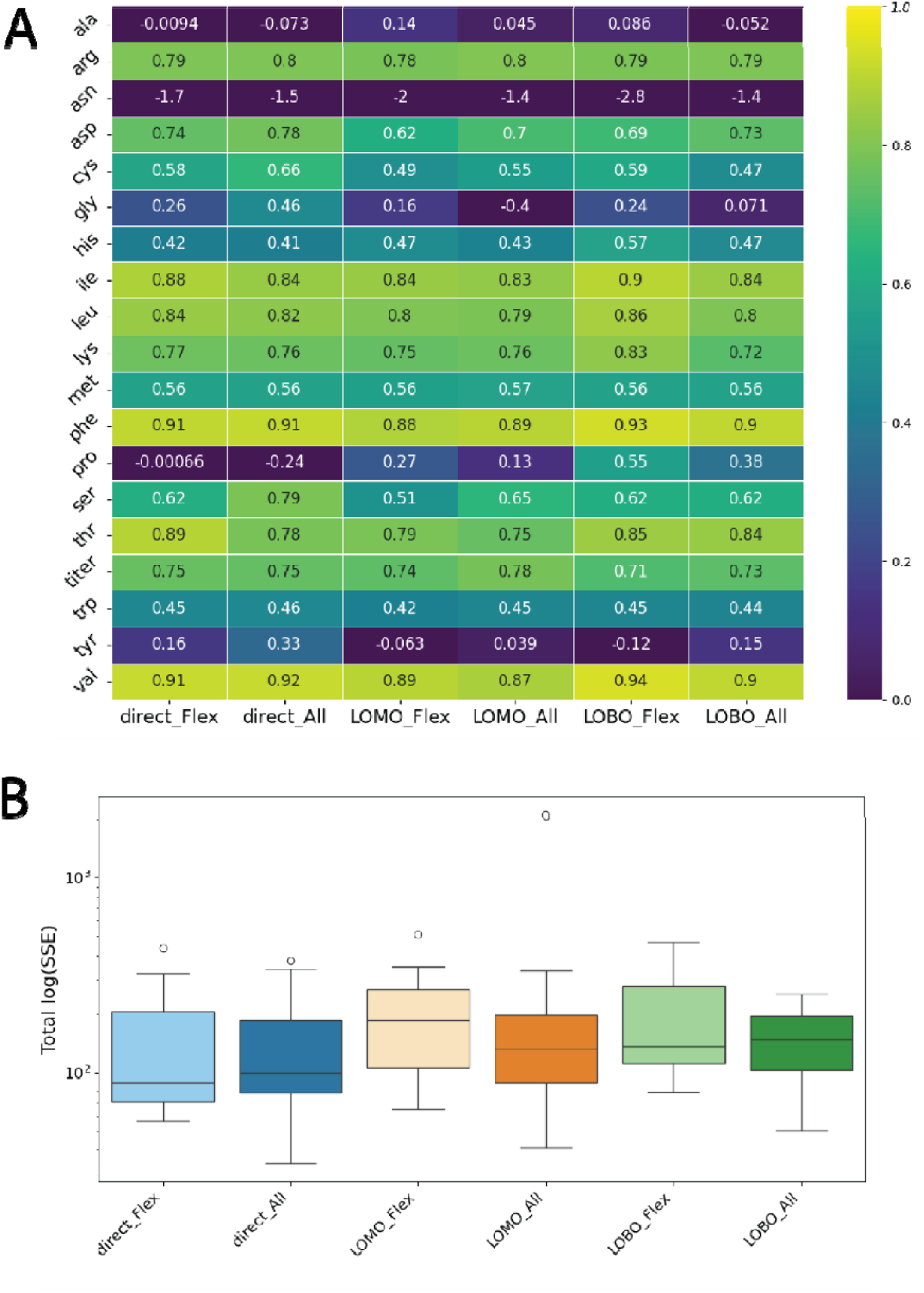
PC-dFBA prediction performance across ANN configurations and validation strategies. (A) Coefficient of determination (R^2^) for 19 non-trivial extracellular metabolites under six ANN x validation combinations (direct, LOMO, and LOBO for both *Model_Flex* and *Model_All*). (B) Total log-transformed summed squared error (log(SSE)) distributions for each ANN and validation types configuration.

Despite the increased stringency of the LOMO and LOBO schemes, the overall predictive performance remained comparable to that obtained under direct validation, confirming that the ANN-based PC-dFBA generalizes well across distinct culture media and batches. The total log-transformed SSE distributions (Figure 5B) further support this conclusion, showing overlapping error ranges across all configurations and validation strategies. Both the *Model_All* and *Model_Flex* variants performed similarly, with only marginal differences observed for specific metabolites. A more detailed breakdown of metabolite-specific errors is provided in Supplementary Figure 6, which displays the log(SSE) distributions across metabolites for each validation scheme (top panel) and model configuration (bottom panel).

Finally, a comparison against the original PCA-dFBA implementation (Supplementary Figure 7) highlights the advantages of incorporating ANN-based regression to predict the PCA loadings dynamically. While both approaches yield broadly consistent metabolite prediction trends, the original PCA-dFBA, based purely on static PCA projections, tends to outperform the ANN-driven PC-dFBA for certain well-correlated metabolites such as the product titer and several amino acids (e.g., aspartate, phenylalanine, and valine). However, PC-dFBA demonstrates more stable cross-validation performance, maintaining comparable R^2^ values across direct, LOMO, and LOBO strategies, suggesting improved generalization across distinct culture conditions. Thus, although PCA-dFBA can achieve slightly higher R^2^ values for select metabolites, the ANN-based PC-dFBA offers a more flexible and transferable modeling framework that captures nonlinear flux–loading relationships and retains predictive robustness across unseen domains.

### 3.5. Model integration into the cell-process digital twin

The integration of the neural-network VCD model, the ODE-based Flex-metabolite model, and the metabolic PC-dFBA framework establishes a unified predictive system - herein referred to as the “cell-process digital twin”. This hybrid architecture captures the dynamic coupling between cell growth, metabolism, and bioprocess operation.

To evaluate the predictive capacity of the integrated framework, we performed analyses following the same validation strategies as in the previous section (i.e., direct, leave-one-medium-out (LOMO), and leave-one-batch-out (LOBO)), each applied to both the *Model_Flex* and *Model_All* ANN configurations. Importantly, unlike the previous analysis (Figure 5), where experimentally measured growth and metabolite rates were directly supplied to PC-dFBA, the present setup relies entirely on model-predicted quantities. Thus, the digital twin operates in a fully predictive mode: only the initial concentrations and feeding schedule are provided, without access to any future measurements.

Figure 6 summarizes the validation results for the fully integrated digital twin across the six model configurations (direct, LOMO, and LOBO validations applied to both *Model_Flex* and *Model_All*). Overall, the global performance trends are consistent with those obtained in the previous semi-mechanistic PC-dFBA analysis (Figure 5), indicating that coupling the neural-network and ODE components preserves the model’s predictive structure.

**Figure 6.**
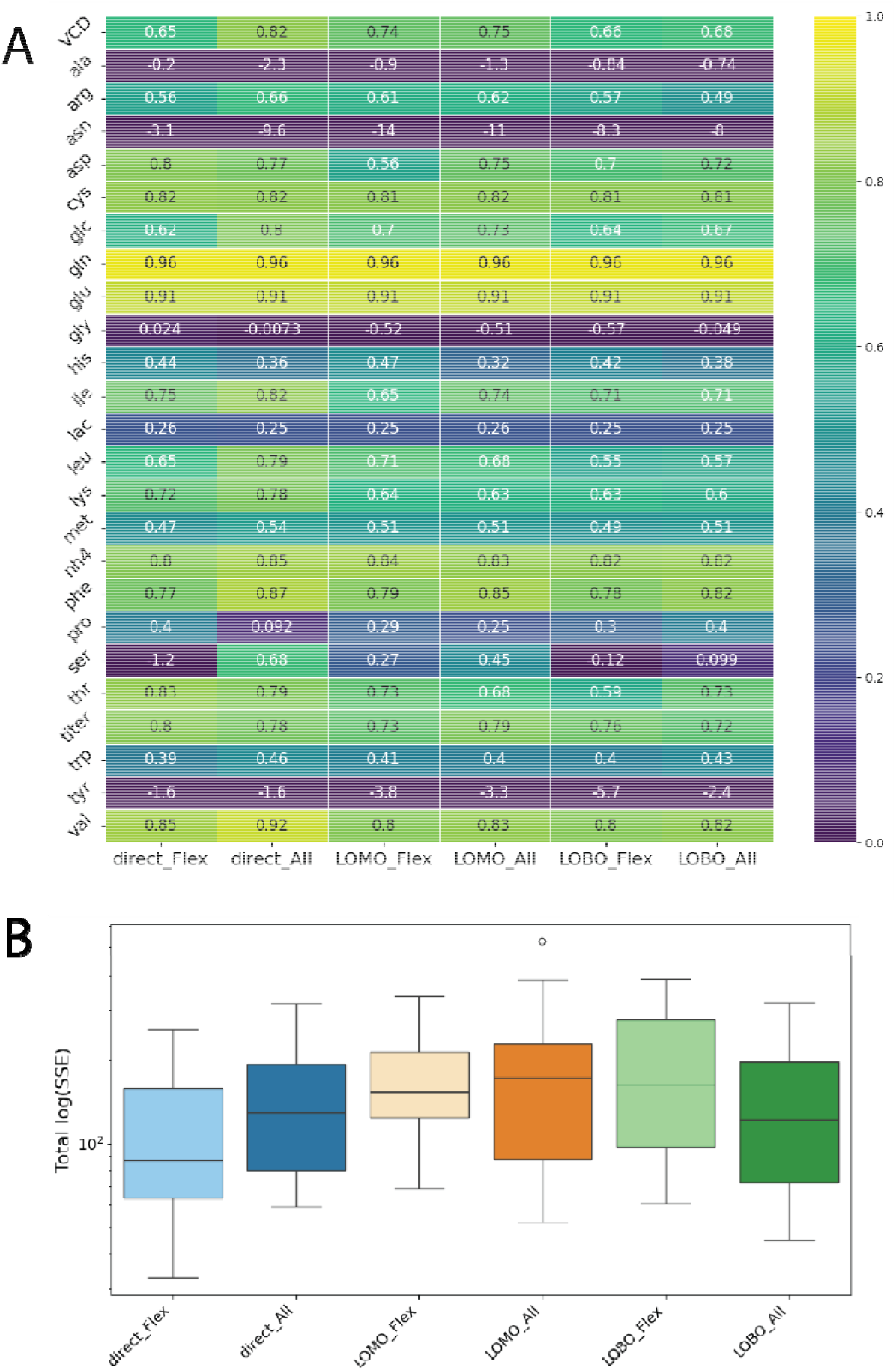
Hybrid modeling framework performance across ANN configurations and validation strategies. (A) Coefficient of determination (R^2^) for 25 extracellular metabolites under six ANN x validation combinations (direct, LOMO, and LOBO for both *Model_Flex* and *Model_All*). (B) Total log-transformed summed squared error (log(SSE)) distributions for each ANN and validation types configuration.

An important aspect of the integration concerns the feedback effect introduced by the neural-network-based VCD predictions. Because the VCD predictor depends on accurate concentration estimates, deviations in those quantities can lead to slight over- or underestimation of the predicted growth rate, which in turn propagates through the ODE and metabolic layers. Indeed, the concentration evolution is dependent on the VCD (the amount of cells defines the variation of metabolite amount, secreted or consumed) but as these concentration data are used to predict the VCD too, there is a “reinforcement loop” that can lead to an increasing model prediction deviation. This “negative-loop” effect can amplify small local prediction errors, particularly in late culture phases. Nevertheless, the overall predictive capability of the full digital twin remains strong, highlighting its resilience to uncertainty propagation and its ability to maintain reliable global process predictions.

As shown in Figure 6B, the “direct x Flex” configuration exhibits the lowest total SSE, confirming that the simpler ANN structure combined with training based on the entire experimental dataset provides the best overall predictive balance. The LOMO and LOBO configurations show comparable predictive performance, suggesting that the framework generalizes robustly even when entire media or individual batches are excluded from training.

Figure 6A further shows that the set of metabolites exhibiting lower predictive accuracy (notably alanine, asparagine, glycine, proline, and tyrosine) remains consistent with the prior results. However, in the integrated framework, serine additionally displays higher variability across validation folds, suggesting increased sensitivity to propagated errors from the growth-rate and metabolite-rate predictions. Despite this, the overall correlation structure remains well preserved, demonstrating that the multi-layer coupling does not degrade performance.

A representative example of such a fully predictive simulation is shown in Figure 7 for batch CS4-5 under the LOMO_Flex configuration. The digital-twin simulation reproduces the temporal evolution of viable-cell density, product titer, and metabolite concentrations with good fidelity. Overall, the results confirm that the integrated digital twin retains the predictive power of its semi-mechanistic components while enabling autonomous, forward simulations from minimal input information.

**Figure 7.**
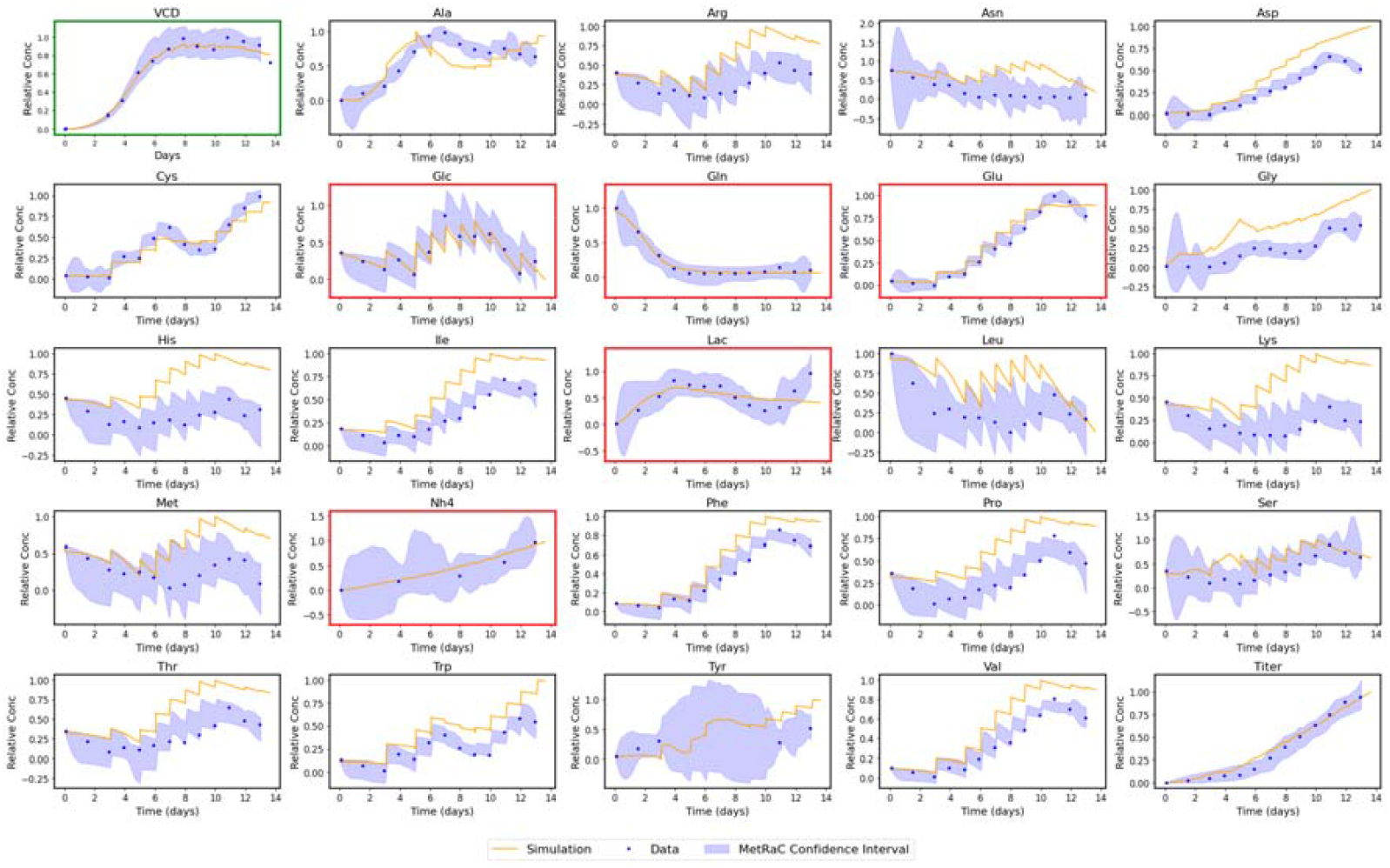
Example of the cell-process digital twin simulation. Comparison of predicted (orange) and experimental (blue) time courses for viable-cell density, product titer, and 23 extracellular metabolites for batch CS4-5 under the LOMO_Flex configuration. The shaded regions represent MetRaC-derived confidence intervals. The green box subplot is associated with the prediction of the VCD NN predictor, and the red box subplots are associated with the prediction of the ODE FLEX model.

## 4. Discussion

This work introduces a cell–process digital twin framework that integrates multiple complementary modeling modules (i.e., machine learning, kinetic ODE-based models, and constraint-based metabolic modeling) to enable fully predictive, dynamic simulation and forecasting of CHO fed-batch bioprocesses. The framework predicts viable cell density, extracellular metabolites, and product titer solely from initial bioprocess conditions and feeding operations, assuming access to a genome-scale metabolic network and prior experimental data. These results demonstrate that mechanistic and machine-learning elements can be synergistically combined to achieve accurate, interpretable, and scalable digital twins for complex mammalian cell culture systems.

Building upon the foundational work of Ramos et al. (2022), we developed the PC-dFBA algorithm, introducing several methodological advances that significantly extend the original dynamic metabolic modeling framework. While the earlier approach established the feasibility of hybrid kinetic–stoichiometric simulation, the present work enhances both the temporal coupling and the data–model integration strategy. First, PC-dFBA implements a fully sequential time-point coupling scheme, ensuring temporal coherence between consecutive simulations and improving the dynamic realism of metabolic flux evolution. Second, it enforces MOMA-based flux continuity constraints, reducing unrealistic discontinuities in predicted flux distributions across time steps. Third, it replaces the static PCA-based loading estimation from Ramos et al. (2022) with a neural-network predictor, enabling adaptive generalization across varying media compositions and process conditions. In parallel, the introduction of a robust median–IQR normalization procedure enhances training stability and facilitates model transferability between datasets of differing scales. Two additional innovations further strengthen the framework: the use of MetRac-based rate estimation, which relies on a Bayesian inference framework to provide more precise and uncertainty-aware kinetic rate estimates, and the implementation of a reduced, data-tailored metabolic network that ensures the model’s structural complexity and feasible solution space remain consistent with the available experimental information. Together, these developments transform the original algorithm into a flexible, modular, and scalable simulation engine, bridging mechanistic interpretability and data-driven adaptability; an essential step toward generalizable, predictive cell-process modeling.

The developed approach holds significant promise for bioprocess development and control. Unlike purely mechanistic models, which require extensive biochemical parameterization, or purely black-box machine-learning approaches, which often lack interpretability, the cell-process digital twin provides a balanced hybrid framework that captures underlying metabolic logic while maintaining adaptability to diverse media compositions and process modes. The framework could be used to develop models for batch, fed-batch, and perfusion operations and is conceptually generalizable to any organism for which a genome-scale metabolic model is available. Consequently, it may be extended beyond CHO systems to microbial, stem-cell, or cell-therapy production platforms. As the biopharmaceutical industry transitions toward data-centric and model-based development, the ability to unify metabolic reasoning with predictive analytics represents a transformative opportunity to accelerate process design, optimization, and control.

Despite these advances, several limitations and gaps remain. The neural-network component used to predict viable cell density introduces a feedback sensitivity that can amplify deviations under certain conditions. Improving this element, potentially through at-line data integration or online retraining, would substantially enhance prediction robustness. Similarly, the ANN-based PCA loading predictor could be redesigned to use more stable and biologically grounded inputs. The current implementation relies on rate-derived features, which may induce sensitivity to local data variability. In the present version, PCA loadings are predicted from the absolute rate profiles of 25 metabolites, and batches with the same media composition share the same PCA loading label. This creates multiple distinct input profiles mapped to a single target, which may reduce the model’s ability to generalize. A more robust approach would be to represent these batches using summary statistics, such as median and interquartile range (IQR) rate values, to define an input range rather than relying on individual rate trajectories. Furthermore, the model does not currently incorporate temporal context (e.g., culture time or a correlated proxy such as accumulated biomaterial), nor does it use biomaterial information directly. Including these temporal and physiological features would provide additional structure to the input space and could significantly improve prediction stability and biological relevance.

Despite the integration of PCA constraints, the global optimization problem underlying PC-dFBA remains underdetermined, implying the existence of non-unique solutions. Extending the framework to a PC-dFVA formulation (i.e., first solving the standard FBA problem, here PC-dFBA, then fixing the optimum obtained with the FBA problem as an additional constraint and after successfully minimizing and maximizing each exchange reaction flux to quantify the variability of the problem under the specified constraints) reveals that the predicted flux variability often overlaps with experimentally observed data ranges, indicating that the model captures biologically plausible and consistent solution spaces. As illustrated in Supplementary Figure 8, this analysis highlights both the strength and the current flexibility of the framework: while the predictions remain coherent with observed trends, the spread of feasible flux distributions suggests that multiple valid trajectories can satisfy the same input conditions. This variability underscores the need for additional constraints or informative data layers (e.g., omics-derived flux ratios, regulatory information, or refined flux-variability bounds) to further restrict the feasible solution space and enhance the specificity and robustness of model predictions. Beyond serving as a validation tool, the PC-dFVA analysis could therefore act as a diagnostic instrument, guiding systematic model refinement and identifying where new data or mechanistic insights would have the greatest impact on improving predictive fidelity.

Several avenues also exist for methodological improvement. The hybrid architecture of PC-dFBA already demonstrates the potential of combining neural-network surrogates with mechanistic equations. Future enhancements could incorporate symbolic regression or automated model-generation tools to identify optimal submodel structures autonomously and reduce expert bias in model formulation. Additionally, physics-informed neural networks (PINNs) could serve as drop-in replacements or complements for ODE-based modules, offering a flexible route to integrate mechanistic priors with data-driven learning. Beyond architectural refinement, another promising direction lies in the systematic identification of key metabolites that should be treated as model inputs rather than outputs to enhance predictive robustness. For instance, metabolites such as alanine could be provided as direct inputs to the PC-dFBA framework to improve dynamic prediction of metabolic interactions. Similarly, while certain metabolites, such as pyruvate, are not currently included among the modeled variables, they are often measured in experimental datasets and could, with appropriate data availability, be integrated into the ODE module to better constrain intracellular flux dynamics. Developing data-driven strategies to identify such informative or complementary metabolites, through sensitivity analysis, feature selection, or model-based relevance ranking, is currently under active investigation. For example, flux coupling can be used to identify correlations between the metabolite predictions, which can be used to cluster metabolites by group/subgroups and identify which one should be used as input to provide enough information about the other from the group. The modular structure of PC-dFBA naturally accommodates these refinements, ensuring that the framework can evolve as new measurements and analytical capabilities become available.

Beyond predictive modeling, the PC-dFBA algorithm opens a broad landscape of opportunities for intelligent bioprocess systems. The current implementation deliberately relies on viable cell density (VCD) and a small panel of five key FLEX metabolites as input variables, precisely because these quantities are routinely measurable at-line in modern bioprocess operations. This design choice was made to ensure that the framework could be realistically extended toward at-line or online deployment, where model predictions and control actions could be continuously updated using readily available process data. In this context, PC-dFBA can function as a soft sensor, providing estimates for unmeasured metabolites or product attributes from these accessible inputs. Such capability could enable adaptive control strategies, where the digital twin dynamically optimizes feed media composition, volume, and timing in real time. Moreover, iterative use of PC-dFBA across production runs would support self-updating digital twins, where each new dataset incrementally refines the model, improving its fidelity and predictive power. This iterative learning paradigm would also facilitate predictive monitoring and early anomaly detection, enabling proactive interventions before process deviations affect product quality. The framework also lends itself to model-based Design of Experiments (DoE), in silico process screening, and Model Predictive Control (MPC) integration for autonomous reactor operation. Embedding PC-dFBA directly into bioreactor control systems could provide near-real-time forecasting, fault detection, and optimization capabilities. By unifying mechanistic transparency with machine-learning flexibility, PC-dFBA establishes a foundation for next-generation self-optimizing bioprocesses.

In summary, PC-dFBA represents a significant step toward the realization of intelligent, adaptive digital twins for biomanufacturing. It bridges the gap between mechanistic process understanding and data-driven prediction, achieving generalizable performance across diverse culture conditions without requiring media-specific model re-engineering. As richer datasets, advanced sensing technologies, and automated metabolic modeling tools continue to emerge, frameworks such as PC-dFBA will play a central role in the evolution toward autonomous, data-informed bioprocess design. Ultimately, these technologies promise to accelerate development timelines, reduce experimental burden, and enhance both product quality and process robustness; key objectives in the digital transformation of biopharmaceutical manufacturing.

## Acknowledgments

We thank Ali Safari and Dirk Muller for providing the experimental data, and the Sartorius Corporate Research team for their support in developing this study.

## Author contributions

A.R. designed and coordinated the study, generated the results, and wrote the manuscript. A.R., A.V., and S.P. developed the data management framework to support the implementation of the algorithm. A.V., A.R., and D.A. developed the methodology related to the calculation of metabolic rates from concentration data. A.V., A.R., and J.J. developed the methodology to predict VCD using a recurrent neural network. A.R. developed the methodology related to the kinetic modeling part of the study. A.A. and A.R. developed the methodology related to the metabolic modeling part of this study. A.R. and J.T. supervised the project execution. All authors reviewed the manuscript.

