## Supplementary Figures - Tables for "A Hybrid Modeling Framework for Predictive Digital Twins of CHO Cell Culture"

**Supplementary Figure 1**. Relative metabolite concentration profiles over time for 23 fed-batch CHO-S cell cultures. The colors in the figure represent the eight distinct FMA+FMB formulations tested. For cultures with the same formulation, the volumes of FMA+FMB added varied, while the timing of additions was consistent across all experiments.

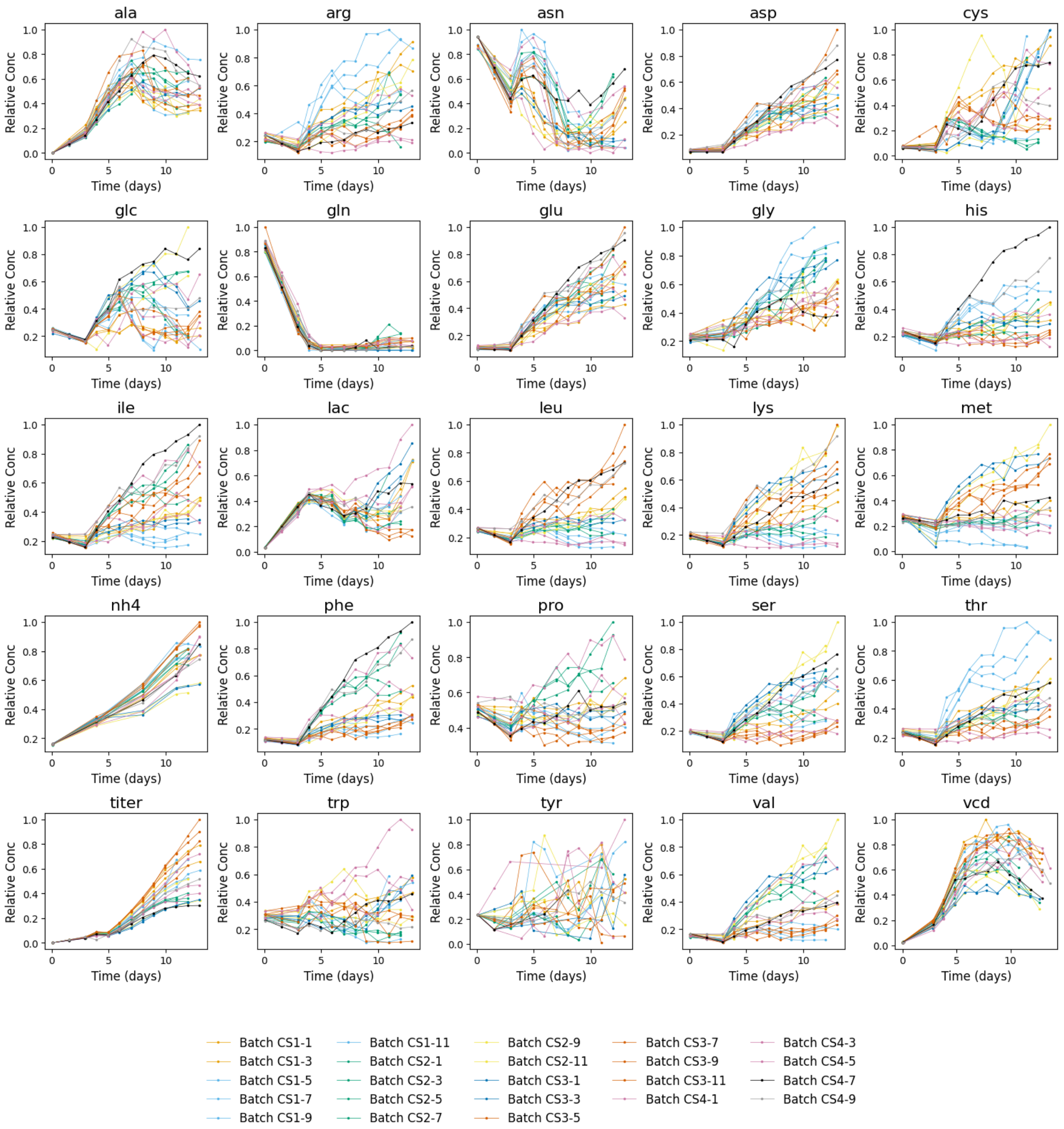

**Supplementary Figure 2**. Schematic representation of the recurrent neural network (RNN) architecture used for growth rate prediction.

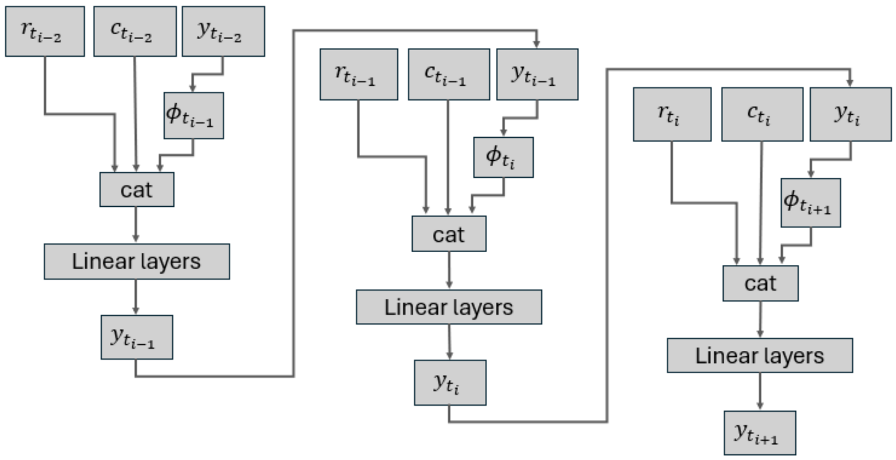

**Supplementary Figure 3.** Comparison between simulated and experimental viable cell density (VCD) trajectories obtained by integrating the NN-predicted growth rates within the ODE framework. The shaded regions denote the MetRac confidence intervals.

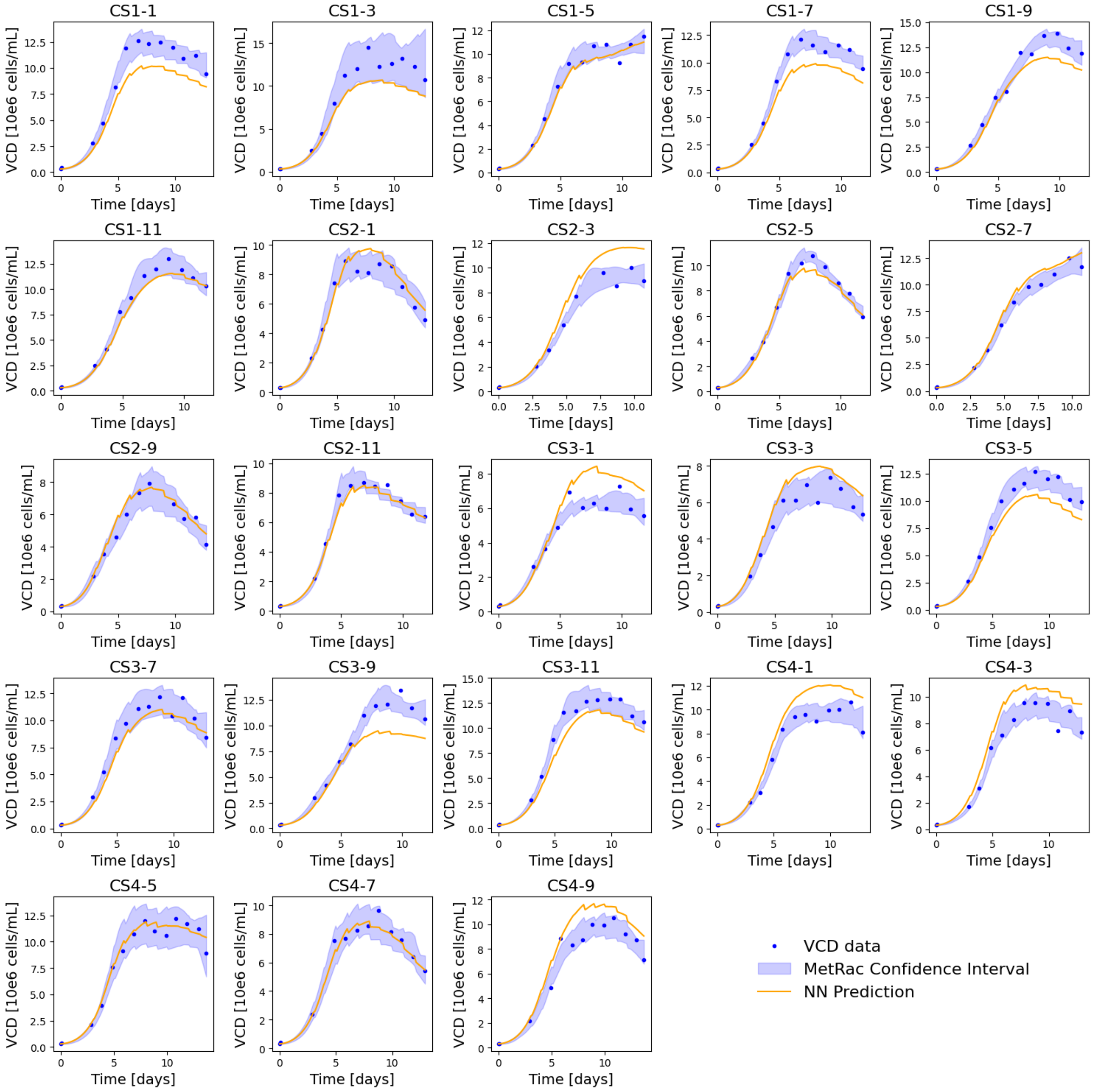

**Supplementary Figure 4.** Overview of model fits and simulated state variables for all batches. Each row corresponds to one batch. Columns 1 and 2 show the fitted model simulations (solid lines) and experimental data (points) for viable and dead cell concentrations, respectively. Columns 3 and 4 display the simulated trajectories of lysed cells and biomaterial (metabolic by-products), which are internal state variables not directly measured experimentally.

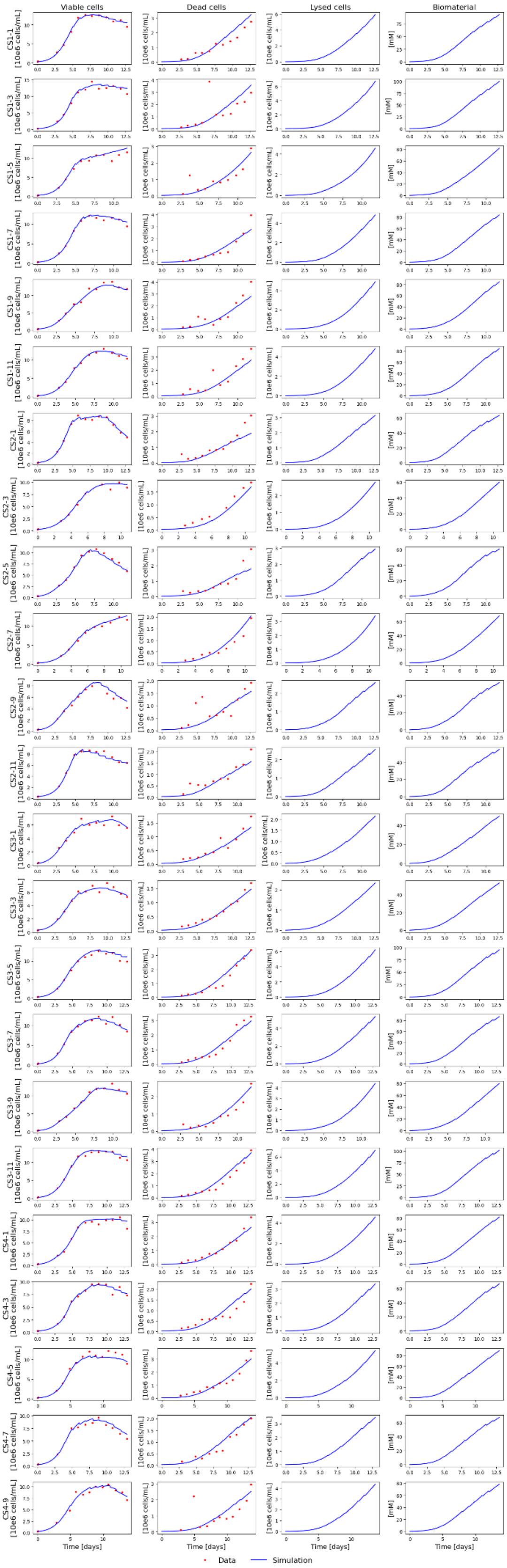

**Supplementary Figure 5.** Detailed metabolite time-course simulations for all batches using the FLEX ODE model. Comparison between experimental data (blue dots), simulation results (orange lines), and confidence intervals (shaded regions) for the five FLEX metabolites: glucose, lactate, glutamine, glutamate, and ammonia. The experimental and simulated concentration data for glucose, lactate, glutamine, glutamate, and ammonia are normalized to be displayed on a relative scale.

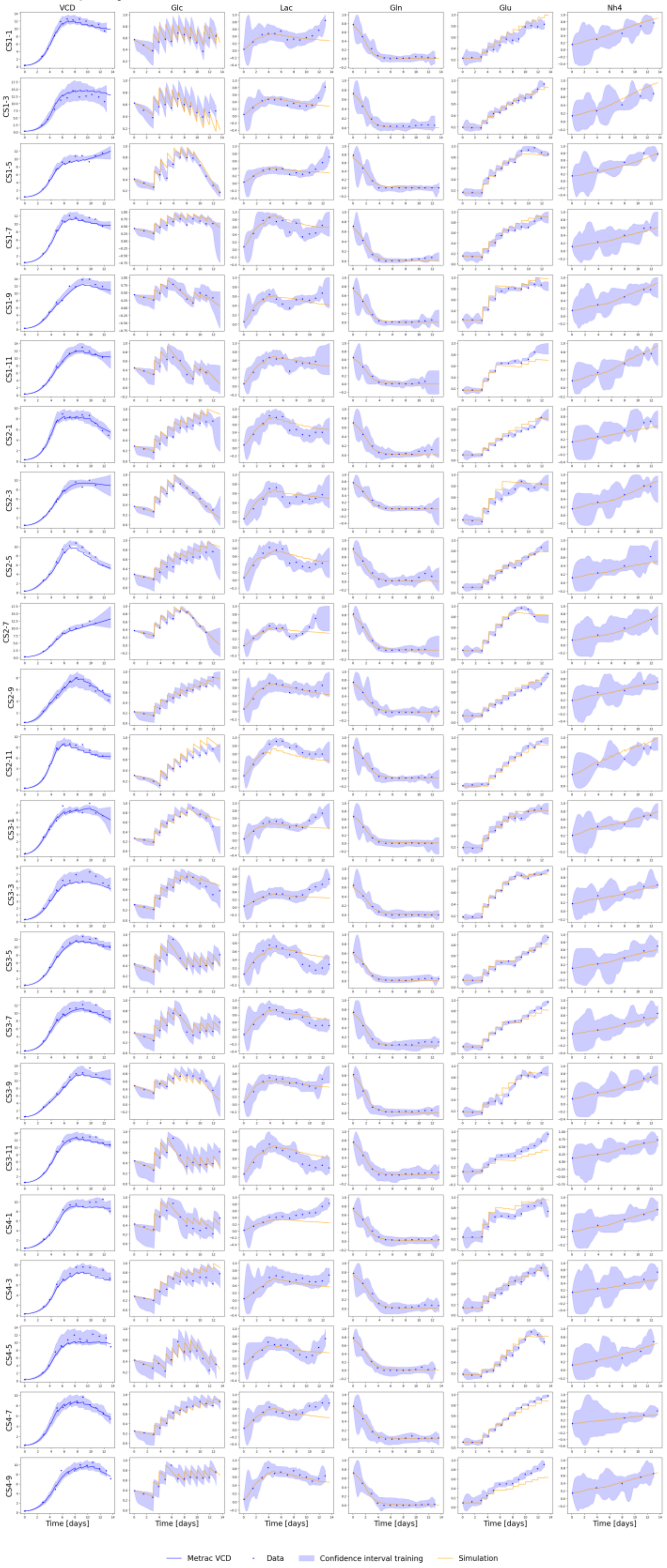

**Supplementary Figure 6.** Metabolite-wise prediction errors for PC-dFBA models. Top panel: Boxplots of log-transformed summed squared errors (log(SSE)) for each metabolite under the three validation strategies (direct, LOMO, LOBO). Bottom panel: Comparison of log(SSE) distributions between Model_Flex and Model_All across metabolites.

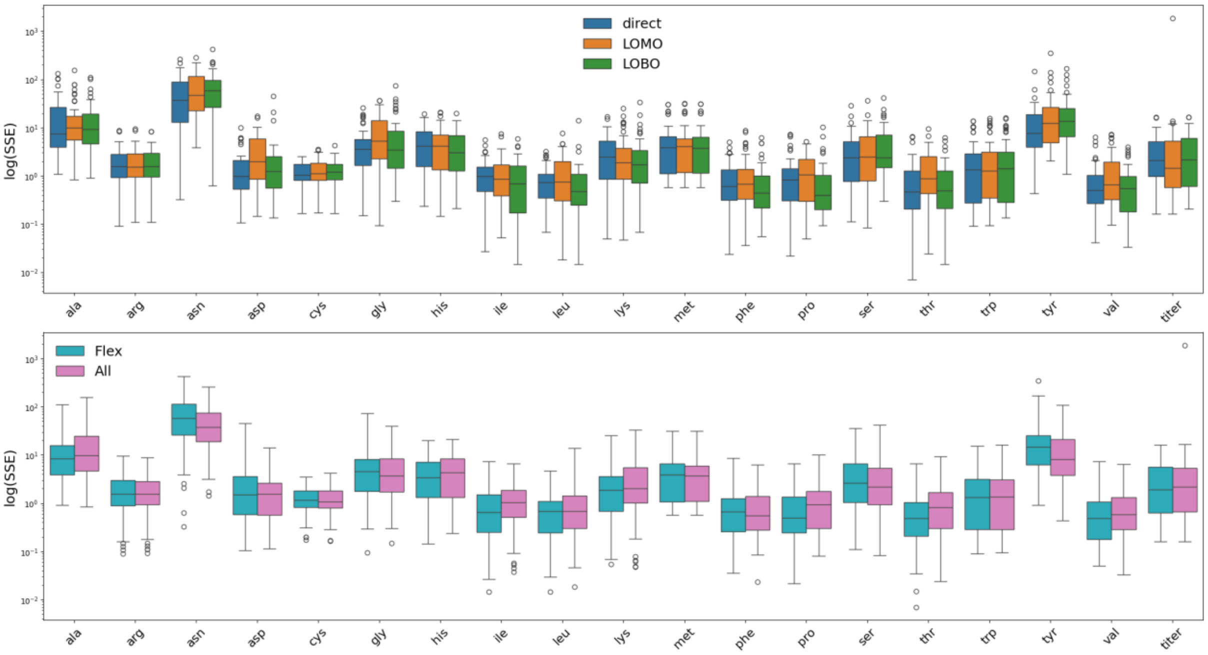

**Supplementary Figure 7.** PCA-dFBA prediction performance across validation strategies. (A) Coefficient of determination ($R^2$) for 19 extracellular metabolites obtained using the original PCA-dFBA formulation across the three validation strategies (direct, LOMO, and LOBO). (B) Total log-transformed summed squared error (log(SSE)) for the same simulations.

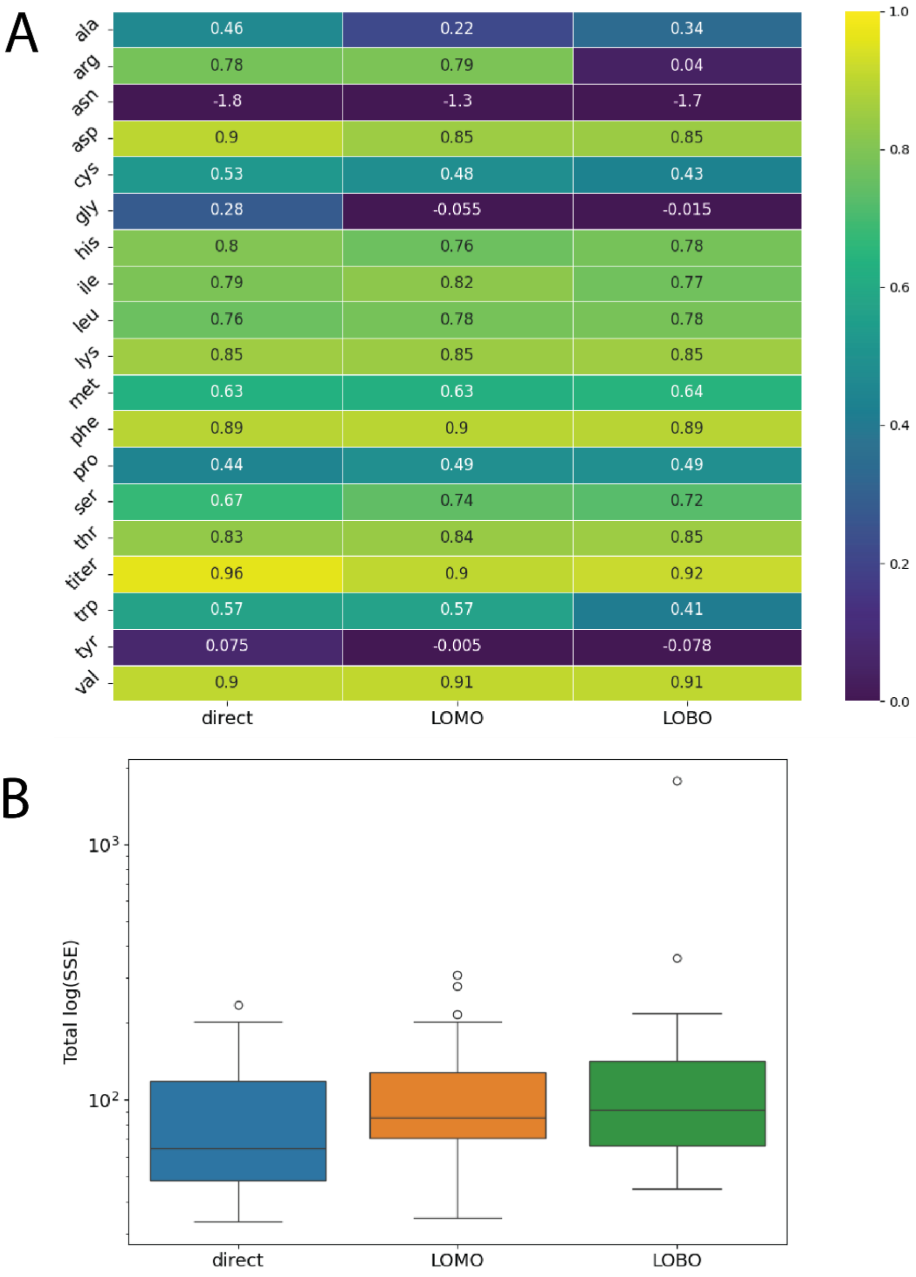

**Supplementary Figure 8. Example of the propagation of the prediction variability of the PC-dFBA algorithm.** Comparison of predicted (orange) and experimental (blue) time courses for viable-cell density, product titer, and 23 extracellular metabolites for batch CS4-5 under the LOMO_Flex configuration. The shaded blue regions represent MetRaC-derived confidence intervals. The shaded green regions represent FVA simulation envelope.

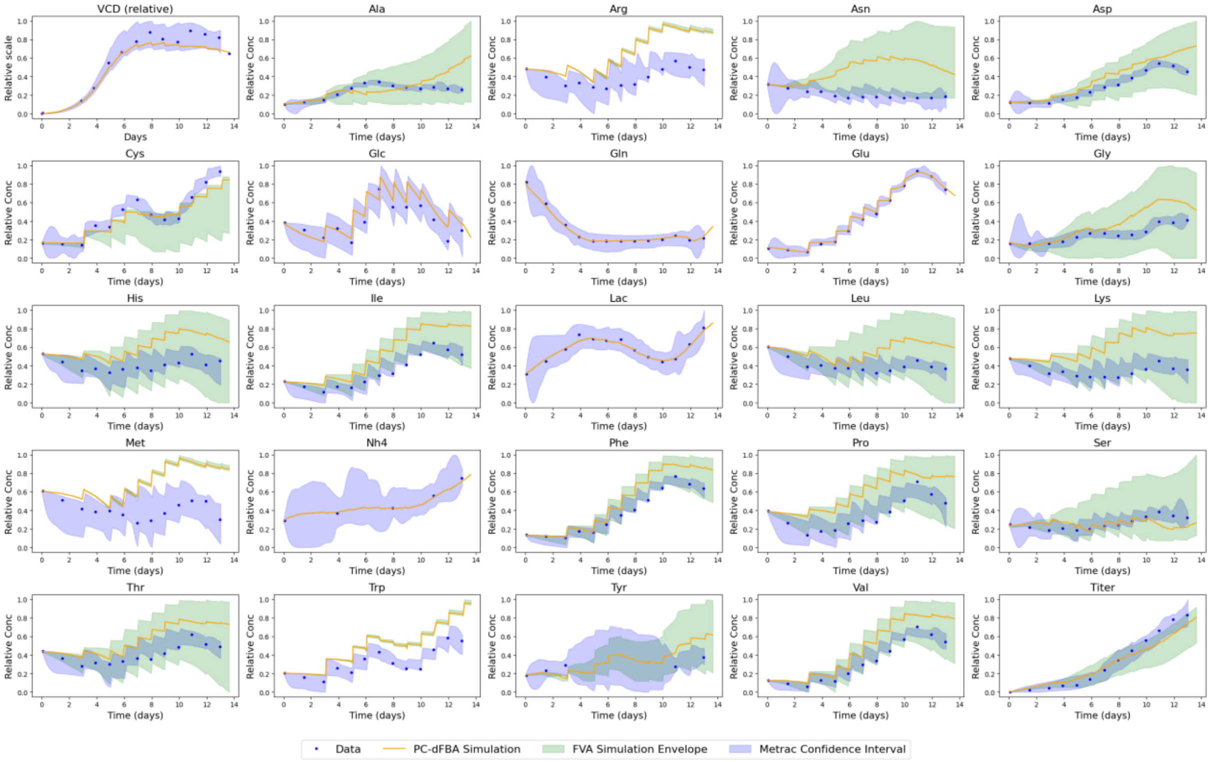

**Supplementary Table 1**. Summary of structural and functional changes in the iCHO1766 model during the five-step reduction process using MetRaC rates (95% confidence interval)

| Model Reduction Steps | Reactions | Metabolites | Exchange Reactions | Genes | Essential Genes |
|---|---|---|---|---|---|
| Initial Model | 6663 | 4455 | 602 | 1766 | 233 |
| Step 1: Resolve Infeasibilities | 4329 | 2231 | 284 | 1492 | 211 |
| Step 2: MILP Exchanges | 3563 | 1797 | 33 | 1366 | 195 |
| Step 3: Transport | 2164 | 1405 | 33 | 1134 | 185 |
| Step 4: pFBA | 575 | 468 | 32 | 595 | 138 |
| Step 5: Thermodynamic Infeasible Loops | 575 | 468 | 32 | 595 | 138 |

**Supplementary Table 2**. Summary of algorithm features evolution from FBA to PC-dFBA

| Version | Key Feature | Temporal | Empirical Constraint Source |
|---|---|---|---|
| FBA | Mechanistic only | Pseudo steady-state | None |
| Hybrid FBA | Adds PCA-based constraints | Pseudo steady-state | PCA loadings |
| Hybrid dFBA | Dynamic extension | Time-discretized | PCA per interval |
| Hybrid dFBA - MOMA | Smooth transitions | Time-discretized | PCA per interval |
| PC-dFBA | ANN-predicted loadings to remove time dependency | Dynamic | NN regression (no interval definition) |
