## Supplementary Methods for "A Hybrid Modeling Framework for Predictive Digital Twins of CHO Cell Culture"

### Evolution of the HybridFBA to PC-dFBA Algorithm

Standard FBA formulates the metabolic network as a linear programming (LP) problem to compute intracellular fluxes under a pre-defined objective function, and a limited set of fundamental assumptions: (i) balanced intracellular metabolite pools (pseudo steady-state hypothesis), (ii) reaction stoichiometry constraints, and (iii) flux irreversibility constraints (i.e., thermodynamic reaction directionality). Together, these represent the "mechanistic" or "parametric" constraints (Orth et al., 2010). The formulation is:

$$max\, J = c^T.v \qquad (S1)$$

subject to

$$S.v = 0 \qquad (S2)$$

$$lb \ \leq \ v \ \leq \ ub \qquad (S3)$$

where $S$ is the stoichiometric matrix, $v$ the vector of fluxes, and $c$ the objective coefficient vector (e.g., maximizing growth).

Ramos *et al.* (2022) introduced HybridFBA, a refinement combining mechanistic (parametric) constraints with empirical (non-parametric) ones derived from Principal Component Analysis (PCA) of experimental flux data. This hybrid formulation extends standard FBA by adding PCA-derived flux correlation constraints and introducing both mechanistic variables ($v$) and empirical variables ($Score$) - the principal component scores - to enforce relationships between the temporal evolution of different metabolites. Thus, HybridFBA simultaneously optimizes $v$ and $Score$ in a single LP problem.

Additional mechanistic constraints are applied only to a subset of the measured exchange fluxes - specifically, the metabolites selected as inputs for the simulation. These fluxes are referred to as "hard bounds" ($v_{HB}$):

$$lb_{HB} \ \leq v_{HB} \leq ub_{HB} \qquad (S4)$$

where $lb_{HB}$ and $ub_{HB}$ corresponds to the lower and upper bounds imposed on these fluxes.

The PCA-derived constraints can be expressed as a set of linear equations that relate the PCA scores to the exchange fluxes (i.e., measured rates) through the PCA loadings.

The PCA is applied to the data matrix of calculated exchange rates (obtained from MetRaC for this study). Data are autoscaled (zero mean, unit variance) to ensure that each metabolite contributes equally. For each principal component $i$:

$$Score_i = \sum_{j=1}^{m} Load_{ij}.\overline{v_{meas,j}} \qquad (S5)$$

where $Load_{ij}$ is the loading of the $j$-th exchange flux on the $i$-th PC, $\overline{v_{meas,j}}$ the normalized (autoscale) measured rate for the metabolite $j$ and $m$ the number of measured metabolites.

The PCA constraints, which combine mechanistic and empirical relationships, can then be expressed as:

$$\mu_{vmeas} - RF.\sigma_{vmeas} \leq (S_e \quad -\sigma_{vmeas} \circ Load\,)\begin{pmatrix} v \\ Score \end{pmatrix} \leq \mu_{vmeas} + RF.\sigma_{vmeas} \qquad (S6)$$

where $S_e$ is the stoichiometric matrix associated with the measured extracellular metabolites, $\mu_{vmeas}$ is the mean of the calculated rate across all data available, $\sigma_{vmeas}$ is its associated standard deviation (the same values used to autoscale the PCA data). ° denotes the Hadamard (element-wise) product and $RF$ is a relaxation factor that controls the allowed mismatch between empirical and model-predicted exchange rates.

An additional inequality constrains the principal component scores:

$$LB_{Score} \leq Score \leq UB_{Score} \qquad (S7)$$

where $LB_{Score}$ and $UB_{Score}$ are the lower and upper bounds calculated for each principal component score.

The overall objective function of the hybrid LP is formulated as:

$$min_{v,Score}\{J = c_v^T v + c_{Score}^T Score\} \qquad (S8)$$

where $c_v$ selects the FBA objective (typically the growth reaction) and $c_{Score}$ is a vector of ones (to minimize the empirical component).

Given the semiparametric nature of HybridFBA, a calibration of the nonparametric constraints is always required. The most important parameter is the number of principal components (NPC), which defines both the number of optimized scores and the number of columns in the loading matrix. Typically, the NPC is selected to capture at least 85% of the total variance.

The original HybridFBA was implemented as a “one-step strategy”, assuming quasi-steady state over approximately 70h of culture, and optimizing the model for that single time interval without accounting for phenotypic evolution.

To simulate dynamic culture behavior, the method was extended to a dynamic FBA (dFBA) framework. Culture time is divided into discrete intervals $\Delta T$ (e.g., 0.5 days), and optimization is performed at each interval. Concentrations are updated based on the predicted fluxes, enabling simulation of full culture dynamics over time.

When incorporating time-windowed PCA into this framework, the constraint becomes:

$$\mu_{vmeas}^{\Delta T} - RF.\sigma_{vmeas}^{\Delta T} \leq (S_e \quad -\sigma_{vmeas}^{\Delta T} \circ Load^{\ \Delta T}) \begin{pmatrix} v \\ Score \end{pmatrix} \leq \mu_{vmeas}^{\Delta T} + RF.\sigma_{vmeas}^{\Delta T} \qquad (S9)$$

where $\Delta T$ denotes the selected time interval. The interval length $\Delta T$ was optimized empirically; an interval of 0.5 days minimized prediction error.

Performing optimization independently at each time interval and concatenating results into a time series can introduce abrupt, non-physical shifts in predicted fluxes between consecutive time points. To prevent this, MOMA (Minimization of Metabolic Adjustment) (Segrè *et al.*, 2002) was incorporated to minimize changes in predicted fluxes between consecutive time steps:

$$min \sum \left(v_{pred}^{t} - v_{pred}^{t-1}\right)^2 \qquad (S10)$$

For the initial time point, where no prior flux distribution exists, standard FBA with PCA-derived constraints is employed as the baseline solution.

Finally, the algorithm was further refined into a fully data-driven framework, referred to as PC-dFBA, by removing the explicit dependency of the PCA-derived constraints on discrete time intervals. Instead of performing time-windowed PCA, we adopted a regression-based approach in which Artificial Neural Networks (ANNs) were trained to predict the PCA loadings ($\sigma \cdot Load$) directly from exchange rate inputs. In this way, the relationship between metabolite exchange rates and PCA loadings is learned continuously across all time points, rather than being tied to specific time intervals. Each principal component (PC) is associated with its own neural network regressor, allowing flexible and nonlinear mapping between experimental fluxes and the empirical constraints used in the optimization. This ANN-based regression approach effectively replaces the static PCA loadings with adaptive, data-driven equivalents, leading to the final formulation termed PC-dFBA, as presented in the main manuscript.
